# Bub1 kinase acts as a signalling hub for the entire *Cryptococcus neoformans* spindle assembly checkpoint pathway

**DOI:** 10.1101/2022.09.21.508923

**Authors:** Ioanna Leontiou, Thomas Davies, Ivan Clark, Koly Aktar, Ardra Pamburayath Suresh, Maria Alba Abad, Christos Spanos, Kyung-Tae Lee, Yong-Sun Bahn, A. Arockia Jeyaprakash, Kevin G. Hardwick

## Abstract

*Cryptococcus neoformans* (*Cn*) is an important human pathogen and a model system for basidiomycetes. Here we carry out a dissection of its spindle assembly checkpoint (SAC), focusing on Bub1 and Bub3. In many eukaryotes, including humans, *Saccharomyces cerevisiae* and *Schizosaccharomyces pombe*, Bub1 underwent gene duplication, generating paralogues referred to as Bub1 and BubR1 (or Mad3). Bub1 has upstream signalling functions at kinetochores, whilst BubR1/Mad3 is a component of the downstream mitotic checkpoint complex (MCC) that delays anaphase onset until all chromosomes are correctly attached. Here we demonstrate that the single CnBub1 protein carries out all the checkpoint roles of both Bub1 kinase and Mad3/BubR1. Proteomic analysis reveals kinetochore targeting via Spc105^KNL1^ and interactions with all downstream SAC components and effectors (Cdc20 and the anaphase promoting complex/cyclosome). We demonstrate that CnBub1 kinase activity is required to maintain prolonged checkpoint arrest. Thus CnBub1 acts as a SAC signalling hub and is a future target for anti-mitotic drugs.

## Introduction

Mitosis is the key step in cell division where cells segregate the two copies of the genome they produced in S phase to their daughter cells. Most cells carry out this segregation with remarkably high fidelity and mitotic control pathways have evolved to ensure that this is the case. One of these is the spindle assembly checkpoint (SAC) which is a mitotic surveillance system^1,2^. It monitors interactions between chromosomes and spindle microtubules and if any attachment errors are apparent it delays the metaphase to anaphase transition. This provides the cell with additional time to carry out error-correction, whereby inappropriate kinetochore-microtubule interactions are disrupted, through the action of Aurora B kinase, the enzymatic core of the chromosomal passenger complex (CPC)^3^. The SAC consists of the Mad and Bub proteins and Mps1 kinase, all of which are highly conserved from yeast to human^2^.

Bub1 was one of the first checkpoint components identified, in the original *bub* (budding uninhibited by benzimidazole) screen for mitotic checkpoint mutants^4^. Intriguingly, Bub1 has been duplicated at least 16 times through evolution, and the two resulting genes usually sub-specialised to have very distinct roles^5^. Bub1 acts as an upstream signalling scaffold: it stably associates with unattached kinetochores, where it gets phosphorylated by Mps1 kinase and then interacts with the Mad1-Mad2 complex^6^. The Bub1-Mad1-Mad2 complex then acts as the catalytic platform for production of Mad2-Cdc20, and thereby the mitotic checkpoint complex (MCC)^7,8^. The second gene (BubR1/Mad3) has a downstream role, directly interacting with Cdc20 and Mad2 as part of the MCC complex (Mad2-Cdc20-BubR1-Bub3)^9,10^ that binds and inhibits Cdc20-APC/C^11,12^. Inhibition of this mitotic E3 ubiquitin ligase stabilises securin and cyclin, thereby delaying the metaphase-to-anaphase transition.

A few organisms, such as *Dictyostelium discoideum, Neurospora crassa*, and *Cryptococcus neoformans* did not duplicate the Bub1 gene^5^. Their Bub1 protein is thought to produce a single polypeptide (sometimes referred to as a MadBub) carrying out all the roles of Bub1 kinase and Mad3/BubR1, but this has yet to be thoroughly tested. Here we analyse in detail the multiple domains and functions of Bub1 in *C. neoformans*, which is an environmental basidiomycete yeast and human fungal pathogen. Although widespread in the global environment, it rarely affects humans as it is successfully cleared by the immune system^13^. However, immunocompromised patients can succumb to infections with the yeast passing from the lung, through the blood-brain barrier, to cause meningitis and death. Globally, cryptococcal disease accounts for almost 20% of deaths in AIDS patients: 152,000 annual cases of *Cryptococcal* meningitis worldwide, mainly in Sub-Saharan Africa, lead to ~112,000 deaths in AIDS patients^14^.

During human infection *Cryptococcus* cells can undergo a striking morphological transition and increase in cell size, forming thick-walled, polyploid Titan cells, 10-100μm in diameter^15^. Titan cells are important as host immune macrophages struggle to engulf them during infections^16^. Remarkably, once formed these Titan cells are then able to undergo a reductive division and successfully bud off small, haploid daughter cells^17^. Most are viable, yet some level of chromosome mis-segregation occurs, leading to aneuploidy^17^. This increases genetic diversity of the fungal population and may on occasions lead to drug-resistance in the clinic^18,19^.

Few studies of *Cryptococcus* mitosis have been reported, but these hint at several fascinating aspects of mitotic regulation in this organism. Its centromeres are regional, contain many full-length and truncated transposable elements silenced by RNAi^20,21^ and are enriched for 5mC DNA methylation^22^ and H3K9me2 marked heterochromatin, although the latter is not mediated by RNAi^20^. Unlike *S.cerevisiae* and *S.pombe*, kinetochores are not clustered in interphase, but they do mature and cluster after the microtubule-organising complexes have clustered to form a bi-polar spindle^23^. Whilst much of the *Cryptococcus* kinetochore is well conserved, proteomics identified a fascinating linker protein, Bridgin, which replaces the whole of the CCAN network^24^. Mitosis is semi-open, enabling another level of cell cycle control on key mitotic regulators, such as Aurora B kinase (Ipl1)^25^. Recently several cyclins and *cdc2* homologues were described in *Cryptococcus*, and Cln1 was shown to control polyploid Titan cell formation^26^. Almost nothing is known about the molecular mechanism of the SAC in this organism, nor its relevance to Titan cell formation or their reductive division to produce haploid daughter cells.

Here we demonstrate that CnBub1 carries out multiple SAC functions: site-specific loss-of-function phenotypes, physical interactions, and *in vivo* behaviour are all consistent with it carrying out checkpoint roles of both Bub1 kinase and Mad3/BubR1. Thus it acts as a signalling hub, directly linking kinetochores (Spc105) to APC/C regulation (see Fig.1a). In addition, we describe a novel microfluidics assay for high-throughput analysis of single-cell SAC arrests, which will be transferable to many other stresses and systems.

**Figure 1.**
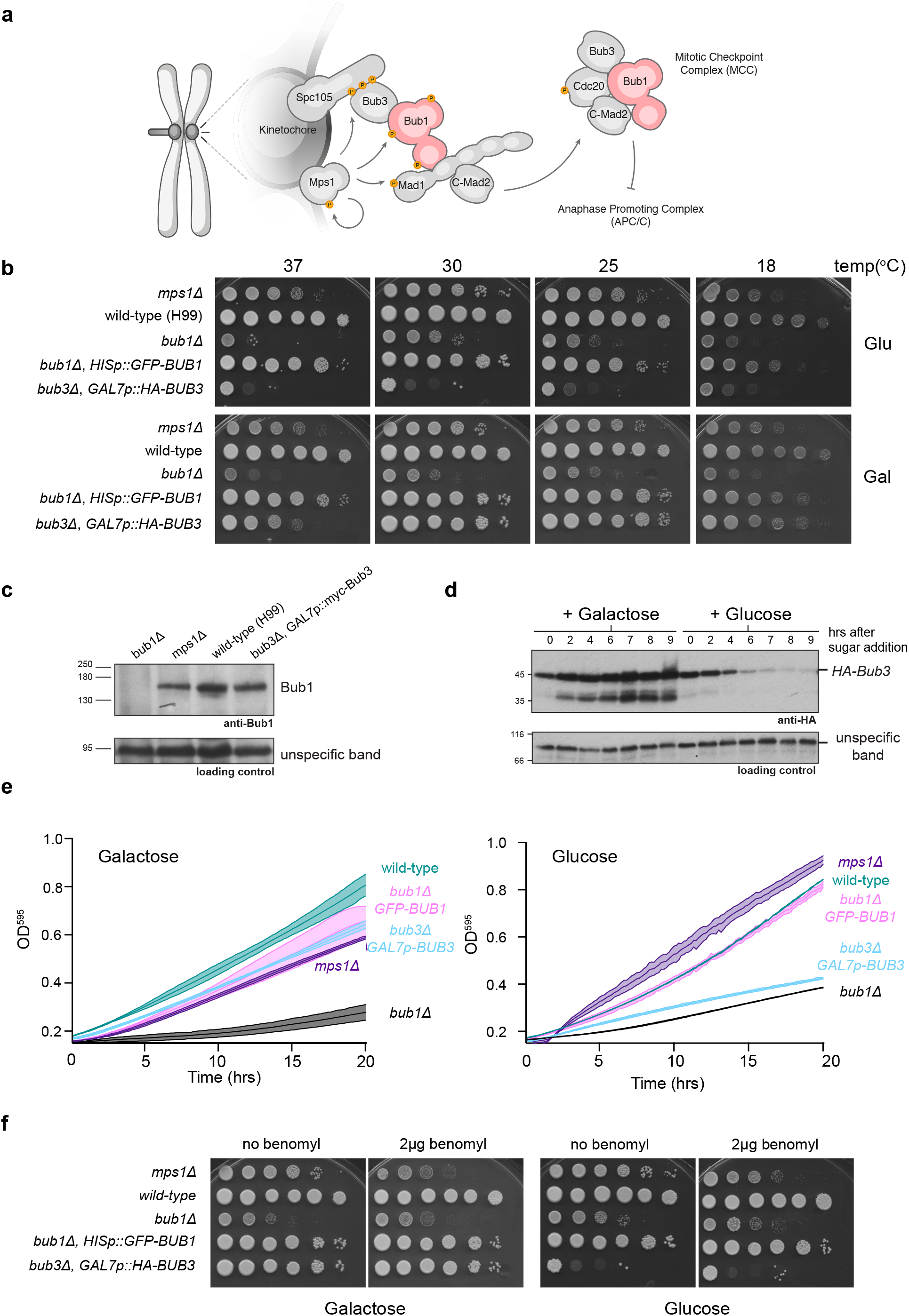
Bub1 and Bub3 are required for normal growth. a) Schematic model of the SAC in *Cryptococcus neoformans*. This is based on previous work from model systems and human cells. The unattached kinetochore recruits active Mps1 kinase. This phosphorylates Spc105 on conserved MELT motifs, to produce a binding site for Bub3-Bub1 complexes. Bub1 is then phosphorylated to produce a binding site for the Mad1-Mad2 complex. The Bub1-Mad1 scaffold provides a scaffold for the efficient assembly of the mitotic checkpoint complex (MCC), which will then bind and inhibit the anaphase promoting complex (cyclosome) APC/C. b) The *bub* mutants grow slowly at all temperatures, in particular 37 and 18°C. Strains were diluted, plated and grown for 2-3 days. The *bub* mutants are both rescued with constructs expressing tagged, but otherwise wild-type copies of the gene from ectopic, safe-haven loci (*P*_GAL7_*HA-BUB3* and *P*_HIS_*GFP-BUB1*). *mps1Δ* is included here as a checkpoint mutant control. c) Whole yeast extracts were made, separated by SDS-PAGE and immunoblotted with polyclonal anti-Bub1 antibodies. d) To determine how quickly HA-Bub3 was depleted from cells, we carried out a time course experiment where log phase cells expressing *P*_GAL7_*HA-BUB3* were pregrown in galactose media, washed 4 times, and then grown for a further 9 hours in either 2% galactose or 2% glucose media. Whole cell lysates were made and immunoblotted with 12CA5 (anti-HA) monoclonal antibodies. e) The strains indicated were grown in Tecan plate readers at 30°C overnight. f) The strains indicated were diluted, plated and grown on YPGlucose or YPGalactose plates with or without 2μg benomyl (anti-microtubule drug) for 3 days at 30°C.

## Results

### *Cryptococcus bub1Δ* and *bub3Δ* strains are extremely sick

In a previous study a systematic set of gene deletions was generated for 129 protein kinases in *Cryptococcus neoformans*^27^. *CnMPS1* was fully deleted and a (C-terminal) fragment of the *CnBUB1* gene was replaced with the *NAT* marker. Both kinases were shown to be relevant to cryptococcal infections as the deletion mutants displayed reduced virulence^27^. We have since made a complete deletion of the *CnBUB1* gene (SuppFig.1a), along with a new rescue construct (*P_HIS3_ GFP-Bub1*, integrated at a safe-haven locus on Chr3). The full *bub1* deletion strain exhibited slow growth at all temperatures and was extremely sensitive to the microtubule poison benomyl (Fig.1b,e,f).

In most systems Bub1 forms a constitutive complex with Bub3^28^. Our expectation was that the *Cnbub3Δ* phenotype would be as equally severe as *bub1Δ*, so first we introduced a second copy of the *CnBUB3* gene that was N-terminally tagged with the HA-epitope, regulated by the galactose-inducible promoter (*P_GAL7_*)^29^ and integrated at the safe haven locus on Chr3. Next, we transformed this strain with a *bub3* deletion blaster cassette^30^ replacing the endogenous *BUB3* gene with the *amdS* (acetamide) marker, and maintained cells on galactose until knock-out transformants were confirmed by PCR (SuppFig.1b). As expected, switching these cells from galactose to glucose media led to gradual depletion of HA-tagged Bub3 (Fig.1d) and cells with a very similar phenotype to *bub1Δ*. The absence of Bub3 leads to reduced growth at all temperatures tested, and extreme benomyl (anti-microtubule drug) sensitivity (Fig.1b,e,f). Growth in plate readers confirms that *bub1Δ* and *bub3Δ* strains grow very slowly compared to *mps1Δ* and the parental, wild-type (H99) strain (Fig.1e). We conclude that, as in *S. cerevisiae*, the *Cnbub1* and *bub3* deletion mutants are both extremely sick under a broad range of growth conditions.

To confirm the *bub1Δ* and test whether the absence of Bub3 or Mps1 proteins might affect Bub1 protein stability we made specific anti-CnBub1 polyclonal antibodies. We expressed residues 53-209aa of bub1Δs as 6xHis-Bub1 fusion protein in bacteria, purified the recombinant protein and used this as antigen. Affinity-purified sheep serum demonstrates that the Bub1 band is missing in *bub1Δ* extracts, but present at normal levels in both the *bub3Δ* and *mps1Δ* strains (Fig.1c).

### CnBub1 functions are Bub3-dependent

To determine the level of dependency of CnBub1 functions on Bub3, we made a site-directed mutation in Bub1 that should abolish Bub3 binding. In the absence of crystal structures, we generated alpha-fold models for the CnBub1-CnBub3 complex using a ColabFold notebook AlphaFold2_advanced. The high confidence model shown in Fig. 2a suggests that CnBub1 indeed interacts with Bub3 via a region containing what is often referred to as the GLEBS motif due to a similar domain in Gle2^31,32^, in a mode similar to that observed for human and *S.cerevisiae* Bub proteins. Particularly, CnBub1 GLEBS motif residues E448 and E449 make electrostatic interactions with CnBub3 R190 and R138, respectively. To disrupt this interaction, we mutated from CnBub1 448E and 449E to K. If all CnBub1 functions are Bub3-dependent, one would expect a very serious (possibly complete) loss of function phenotype for this mutation, as no Bub1 functions could be effectively performed in the absence of Bub3 binding. *bub1-glebs* has a similar phenotype to *bub1Δ* and *bub3Δ* mutants, in terms of benomyl sensitivity (Fig.2b). Plate reader experiments confirm that the *bub1-glebs* mutant has a very similar growth rate to *bub1Δ* and *bub3Δ* strains (Fig.2c). We conclude that most, possibly all, CnBub1 functions are likely to be Bub3-dependent.

**Figure 2.**
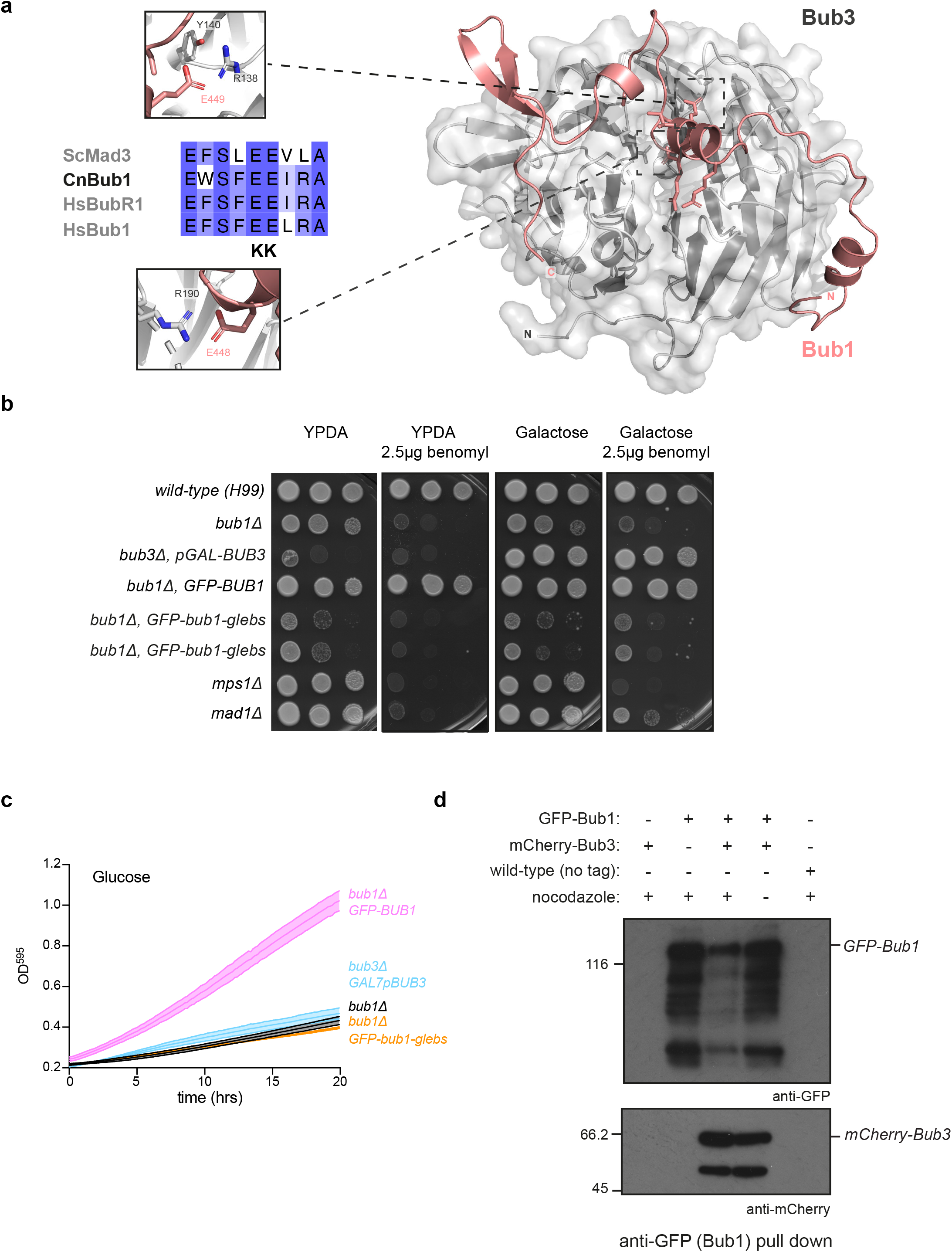
Bub1-Bub3 interactions are critical for Bub1 functions. a) An alphafold model of the Bub1-Bub3 interaction site highlighting the mutated residues. Alignment of the GLEBS motif in *Saccharomyces cerevisiae* Mad3, *Cryptococcus neoformans* Bub1 and human BubR1 and Bub1. In the *bub1-glebs* mutant, resides 448 and 449 were mutated from E to K. b) Sensitivity of *bub1-glebs* mutant. Mutating these two residues E448/9 to K448/9 generates a strain with similar phenotypes to the *bub1Δ* and *bub3Δ* strains. Strains were diluted, plated and grown on YPGlucose or YPGalactose plates with or without 2.5μg benomyl (anti-MT drug) for 3 days at 30°C. c) Plate reader growth curves demonstrate that the *bub1-glebs* mutant grows as slowly as the *bub1Δ* and *bub3Δ* strains at 30°C. d) Co-immunoprecipitation of GFP-Bub1 with mCherry-Bub3. Native protein lysates were made from the strains indicated, incubated with GFP-TRAP beads, beads washed and bound proteins immunoblotted with anti-mCherry and anti-GFP antibodies. mCherry-Bub3 specifically co-immunoprecipitates from strains expressing GFP-Bub1. NB. The difference in Bub1-GFP levels (lanes 2-4) was not reproducible. The faster migrating bands are GFP-Bub1 degradation products,

To demonstrate this interaction biochemically, we co-immunoprecipitated GFP-Bub1 and mCherry-Bub3 from *Cryptococcus* cell extracts. Strains were grown to log phase and some then treated with the anti-microtubule drug nocodazole (2.5μg/ml for 120mins at 30°C) to arrest cells in mitosis. Fig.2d shows that mCherry-Bub3 could be immunoprecipitated with GFP-Bub1 from cycling cell and mitotically arrested cell extracts, consistent with CnBub1-Bub3 being a constitutive protein-protein interaction, throughout the cell cycle.

### CnBub1 co-localises with Bub3 at mitotic kinetochores

In most systems Bub1 targeting to mitotic kinetochore structures is Bub3-dependent^32^. Both Bub proteins are recruited to kinetochores early in mitosis, and in the presence of defective/inappropriate kinetochore-microtubule interactions they are maintained there at high levels. Fig.3a is a schematic of the typical Bub1 recruitment mechanism: when the SAC is activated Mps1 kinase phosphorylates Spc105 on conserved MELT motifs^33–36^, to which the Bub3-Bub1 complex can then bind. Fig.3b shows a field of GFP-Bub1 cells, in the absence and presence of nocodazole. Whilst only the few mitotic cells show foci of GFP-Bub1 in the cycling population, all arrested cells display bright Bub1 foci. To analyse the mitotic cells in more detail GFP-Bub1 was expressed in a strain expressing Tub4-RFP (γ-tubulin) to label the spindle poles^24^. Fig.3c shows GFP-Bub1 in a cycling population of cells, where most cells have diffuse Bub1 staining. Only in the mitotic cells does Bub1 get recruited to structures and these lie between the poles of a bi-polar spindle, consistent with GFP-Bub1 being recruited to mitotic kinetochores. Fig.3d shows a montage of cells taken from a cycling population, highlighting different stages of mitosis. The Bub1-GFP signals are brightest early in mitosis, after the spindle has moved to the bud which is typical of basidiomycetes^23^. After bi-polar spindles form and elongate back into the mother cell the Bub1-GFP signals are significantly weaker. This is consistent with Bub1-GFP being enriched on kinetochores yet to form mature, bi-oriented attachments. In nocodazole-arrested cells the GFP-Bub1 is near to, but distinct from the γ-tubulin (Fig.3e). To confirm that Bub1 is localised to unattached kinetochores we co-localised GFP-Bub1 with the kinetochore marker Dad2 (Fig.3f), which is part of the microtubule-binding Dam1 complex^37 24^. Fig.3g demonstrates that GFP-Bub1 co-localises with mCherry-Bub3 in nocodazole-arrested cells. Finally, Fig.3h shows that GFP-Bub1 is no longer recruited to kinetochores when the Bub3 protein has been depleted from cells, demonstrating that kinetochore targeting of CnBub1 is Bub3-dependent. We conclude that both Bub1 and Bub3 are recruited to *cryptococcus* kinetochores early in mitosis and remain there if kinetochores are unattached.

**Figure 3.**
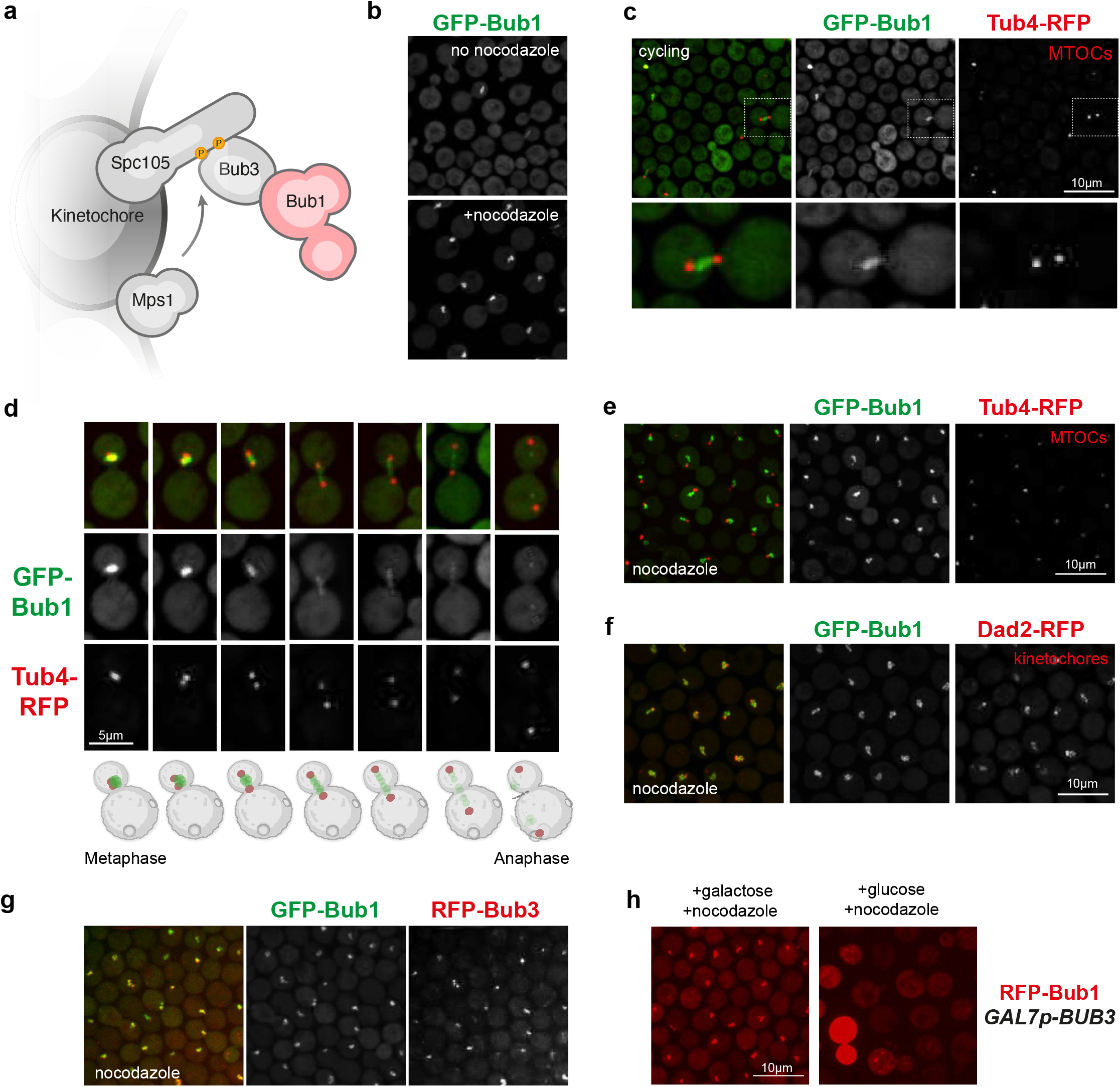
GFP-Bub1 is recruited to mitotic kinetochores. a) Recruitment hypothesis: unattached kinetochores recruit active Mps1 kinase. This phosphorylates Spc105 on conserved MELT motifs, to produce a binding site for Bub3-Bub1 complexes. b) GFP-Bub1 is enriched in bright foci in nocodazole arrested cells. c) Double label imaging of GFP-Bub1 with mCherry-Tub4 (gamma-tubulin, MTOC and spindle pole marker) indicates that in most cells Bub1-GFP is found diffusely through the cell. In mitotic cells with a bi-polar spindle (mid-mitosis) GFP-Bub1 is found between the spindle poles. Scale bars is 10 microns. d) Montage of mitotic cells reveals that GFP-Bub1 is recruited at high levels to kinetochores early in mitosis, when spindles are first being assembled and KTs not yet appropriately attached. Significantly lower levels of GFP-Bub1 are present on kinetochores at later stages in mitosis. Tub4-RFP (gamma-tubulin) is used here as a marker for the spindle poles. These specific images, representative of different stages in mitosis, were taken from a large field of cycling cells. Scale bar is 5 microns. e) Upon nocodazole treatment, bi-polar spindles are not assembled and GFP-Bub1, whilst often nearby is not co-localising with spindle poles (Tub4). f) GFP-Bub1 does co-localise with the kinetochore marker Dad2-RFP in nocodazole arrested (pro-metaphase) cells. g) GFP-Bub1 co-localises with mCherry-Bub3 in nocodazole arrested (pro-metaphase) cells. h) GFP-Bub1 does not localise to kinetochore foci after cells have been depleted on Bub3 (after 6 hours in glucose).

### Distinct Bub1 domains have important yet separable functions

The schematic in Fig.4a indicates conserved motifs that we identified in CnBub1, based on their sequence homology to regions of Bub1 and BubR1/Mad3 in other organisms. For a recent review on Bub1 and BubR1 conserved domains and SLiMs see^38^. Separation-of-function alleles can be a very useful tool when dissecting the multiple functions of a complex signalling molecule like Bub1 kinase. To test whether these domains had their expected functions, we generated a series of *bub1* strains in which the endogenous *BUB1* gene had been deleted and a mutant *bub1* allele is expressed at an ectopic, safe-haven locus on Chr3^30,39^. The *bub1* strains we made include: *bub1-ken1, bub1-ken2*, (both perturbing Cdc20 interactions), *bub1-cd1* (predicted to have no Mad1 binding^6,40^) and a *bub1-kd* (kinase-dead) allele (Fig.4a).

**Figure 4.**
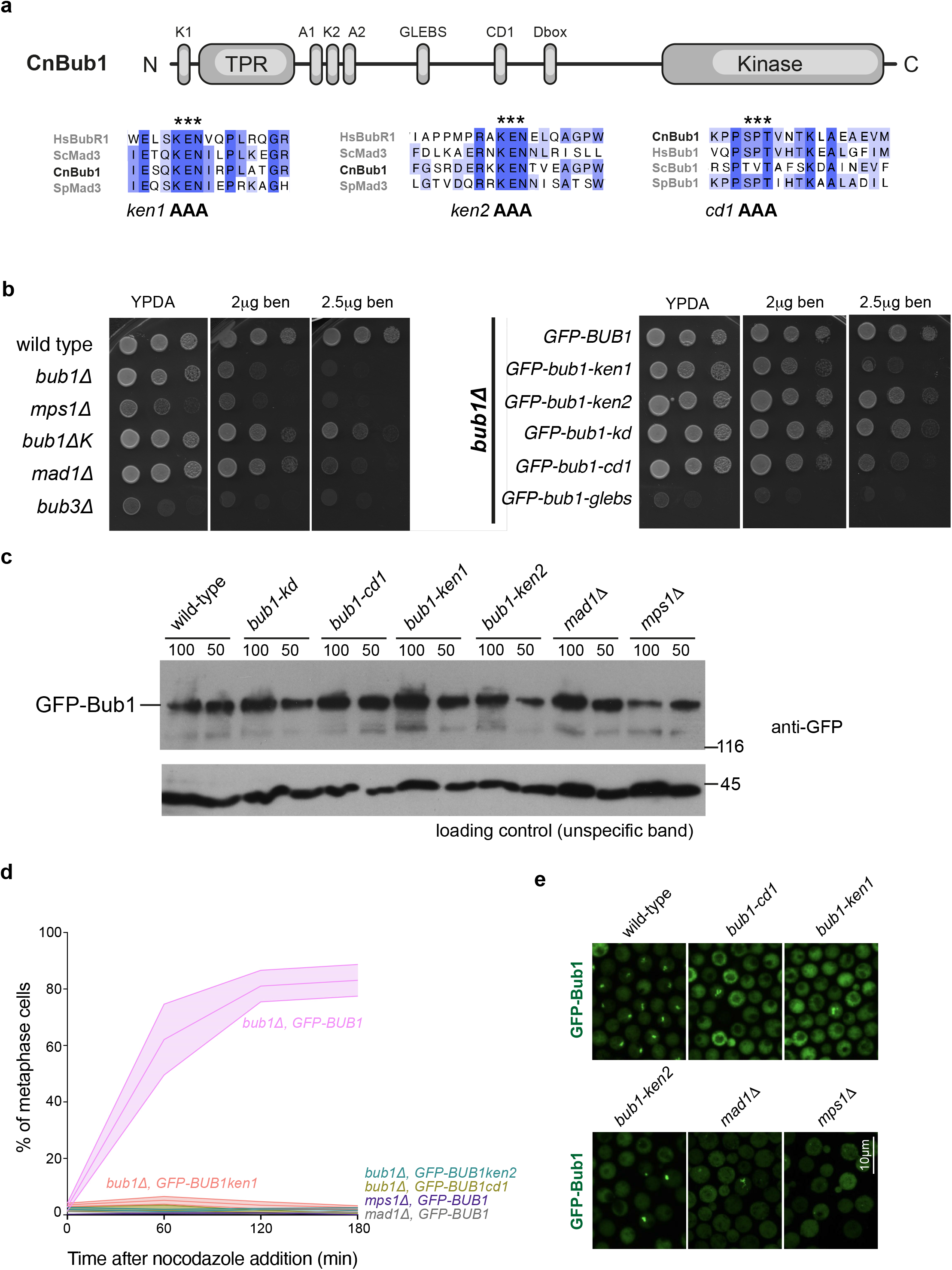
Separation of function alleles, demonstrate distinct functions for Bub1 domains. a) Schematic model of CnBub1 indicating conserved domains, including the two KEN boxes and CD1. Below are Clustal sequence alignments. * indicate the mutations made to generate the *bub1-ken1, bub1-ken2* and *bub1-cd1* alleles. b) The *bub1* alleles, including a kinase-dead (*-kd*) mutant (see Fig. 6) were expressed in a *bub1Δ* background, and phenotypically compared to *mad, bub* and *mps1* deletion mutants. Strains were diluted, plated and grown on YPGlucose (YPDA) plates with 0, 2 or 2.5μg benomyl (anti-MT drug) for 3 days at 30°C. c) Whole cell extracts were made from these strains, separated by SDS-PAGE and immunoblotted with anti-GFP antibodies. All strains are expressing GFP-Bub1 at similar levels to wild-type. The *glebs* mutant protein is less stable (see Supp Fig2). d) The strains indicated, were grown to log phase, then treated with nocodazole at 2.5μg/ml for 60, 120, 180 minutes. They were then imaged and the % of cells with bright GFP-Bub1 foci scored (these represent the mitotically arrested cells). e) Representative images of Bub1-GFP are shown, from the 120min time point. Scale bar is 10 microns.

In an attempt to specifically disrupt spindle checkpoint signalling functions of Bub1, we made mutations in motifs predicted to mediate Cdc20 and Mad1 binding. CnBub1 contains N-terminal motifs (KEN and ABBA motifs) that are typically found in Mad3/BubR1 (Fig. 4a) and very important for interaction with Cdc20^41–43^, and thus formation of MCC and MCC-APC/C complexes. These are key motifs for Mad3/BubR1’s role as a Cdc20-APC/C inhibitor. The CD1 motif (residues 494 to 496 were mutated from SPT to AAA in *bub1-cd1*) is regulated by phosphorylation (typically in a CDK1 and Mps1-dependent fashion). In mitosis this region of Bub1 is phosphorylated and this stabilises a highly conserved interaction with the Mad1 checkpoint protein^6,44,45^. Thus, CD1 is likely to be a key-mediator of Bub1’s checkpoint signalling function. Fig. 4b compares the benomyl sensitivity of these *bub1* alleles with those of *mps1Δ* and *mad1Δ* deletion mutants. The *mad1Δ* mutant will be described in detail elsewhere^46^ and is used here as a checkpoint null mutant. The *bub1-ken1 and bub1-cd1* strains were significantly less sensitive/sick than *bub1Δ*, having a phenotype much more like the *mad* mutant. The *bub1-ken2* and *bub1-kd* strains were less benomyl sensitive, suggesting that their checkpoint responses remain partially intact (see below).

Next we wanted to look at the relative stability of the mutant bub1 proteins. Fig. 4c (and SuppFig2) shows an anti-GFP western blot of these site-specific *bub1* mutants. Only the *bub1-glebs* mutant protein appears to be less stable than wild-type (SuppFig2), so phenotypes of the *bub1-ken, -cd1* and *-kd* mutations are not explained by reduced protein levels. The Bub1 protein was not unstable in the *bub3* mutant, so it is unclear why the *bub1-glebs* mutation is less stable (~50% of wild-type levels, SuppFig2). It is noteworthy that in human cells very low levels of Bub1 protein are sufficient for robust checkpoint signalling and mitotic arrest^47,48^.

There are several reasons why a mutant might be sensitive to anti-microtubule drugs, so more specific checkpoint assays were sought. To assess SAC signalling in *bub1* mutants we challenged cells with the anti-microtubule drug nocodazole and then followed a time course where we scored the numbers of cells in the population that were in early mitosis with bright GFP-Bub1 foci on their unattached kinetochores. If the SAC is proficient then arrested cells should accumulate in the population. Fig. 4d shows fixed time point analysis of the ability of different strains to arrest with Bub1-GFP on kinetochores. All these *bub1* alleles, and the *mad1* and *mps1* mutants, were unable to arrest in mitosis with GFP-Bub1 on kinetochores for a prolonged period. Fig.4e shows that bright GFP-Bub1 foci were detectable in a few of the *bub1* and *mad1* cells, but never in the *mps1* mutant.

The weaker benomyl sensitivity observed in Fig.4b, hinted at a weak SAC response in the *bub1-ken2* and *bub1-kd* alleles. To assess SAC function more carefully we developed a single-cell microfluidics assay, where we could analyse the response of specific cells to microtubule perturbation over several divisions. This was based on the *Alcatras* system, previously developed by the Swain lab^49^. In this assay, cells are pre-grown then trapped as single cells in a microfluidics device where they are maintained in a constant flow of media (see Methods). Over time the trapped cells bud and divide to produce daughters, most of which are washed away by the fluid flow through the device. After 5 hrs in these traps we switched to media containing 2.5 μg/ml nocodazole, and continued our analysis for another 7 hours (30°C), capturing images every 2 minutes. Fig.5a displays temporal heat map representations of 30 randomly selected cells for each strain and Fig.5b a summary plot showing the average dwell time of the enriched GFP-Bub1 fluorescence at the kinetochore within each population. Bright GFP-Bub1 signals represent cells that are in early/mid mitosis. Any mutant that is checkpoint defective will fail to maintain this mitotic delay and thus fail to maintain high levels of GFP-Bub1 on kinetochores through the heat maps. Importantly, strains with a partially functional checkpoint will delay for longer than those completely defective in checkpoint signalling. Thus, this assay provides a quantitative measure of how important specific motifs/domains are in CnBub1 for spindle checkpoint activation and maintenance.

**Figure 5.**
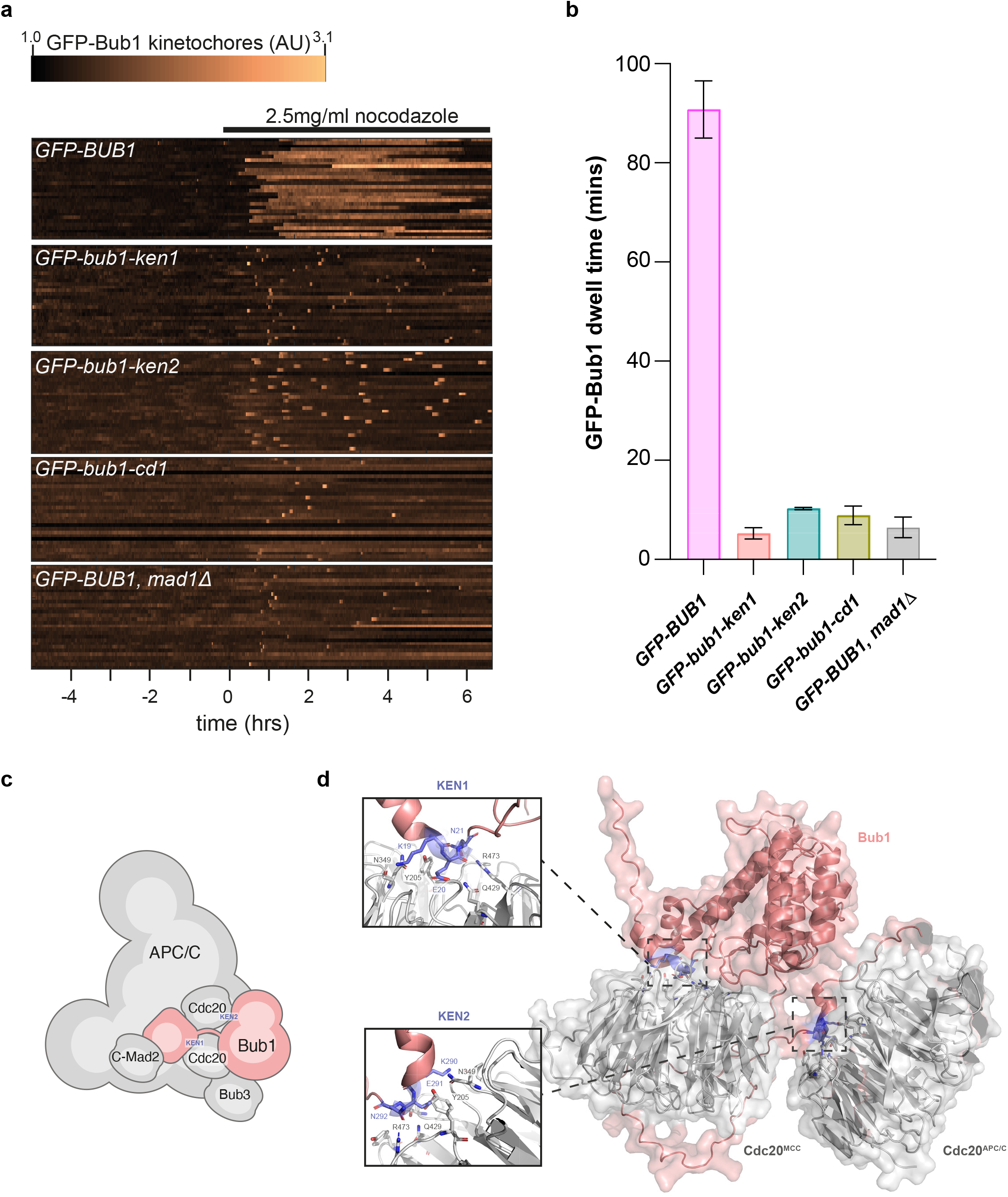
Microfluidics analysis demonstrates varying SAC responses in different *bub1* alleles. a) The strains indicated were cultured in SC media in the microfluidic device for 5 hours, then switched to media containing 2.5μg/ml nocodazole for another 7 hours. Images were captured every 2 minutes. Here 30 random kymographs for GFP- Bub1 are shown for each strain. Bright GFP-Bub1 signals (>1.22) are indicative of mitosis, with significant GFP-Bub1 enrichment on kinetochores. The mutants are unable to maintain mitotic arrests, and the length of the bright ‘dash’ in each heat map indicates how long the cell spent in mitosis. The *mad1D, bub1-ken1* and *bub1-cd1* mutant spend ~5 mins each mitosis, but this is extended slightly in *bub1-ken2* (to ~10 mins). b) Quantitative analysis of GFP aggregation data, shows the average length of time spent in mitosis for each of these strains (n of 386 *BUB1-GFP*; 219 *GFP-bub1-ken1*; 234 *GFP-bub1-ken2*; 128 *GFP-bub1-cd1*; 120 for *GFP-BUB1 in mad1Δ*). c) Schematic explaining Bub1-KEN1 and KEN2 interactions: KEN1 interacts with the Cdc20 molecule that is part of the MCC, whereas KEN2 interacts with the second molecule of Cdc20 that is the APC/C activator. d) Alpha-fold model of Bub1 highlighting the two Cdc20 interactions, through KEN1 and KEN2. Unstructured Bub1 residues (217-271) are hidden in this model.

From this analysis we found that the mitotic (SAC) delay was perturbed in all the mutants we tested, but to varying degrees. The *bub1-ken1* and *bub1-cd1* mutants displayed a very similar response to the *mad1Δ*, which we believe to be fully checkpoint defective. Interestingly, the *bub1-ken2* mutant was able to delay for a little longer (~10 mins), suggesting that a weak checkpoint response is mounted in this strain. Fig.5c shows a schematic model explaining our interpretation of this data, based on MCC-APC/C interactions described in yeast and humans^12,50,51^. An Alpha-fold predicted structure of CnBub1 bound to two molecules of Cdc20 via its two KEN boxes (shown in Fig. 5d) supports our model. KEN1 likely forms a key MCC interaction between CnBub1 and Cdc20: Mad3/BubR1-KEN1 (but not KEN2) interaction is essential for MCC formation in model yeasts and humans. Our interpretation is that the Mad1 and Cdc20 interactions mediated by CD1 and KEN1 in CnBub1 are both critical for checkpoint signalling. As in *S.cerevisiae* Mad3, *S.pombe* Mad3 and human BubR1 we see a weaker phenotype for the *ken2* allele^41,52^. KEN2 mediates the Mad3/Bub1 interaction with a second molecule of Cdc20, which acts as the APC/C activator in MCC-APC/C complexes. In the absence of this second Cdc20 interaction, some inhibition of Cdc20 is still possible through Cdc20 sequestration in MCC complexes (Bub3-Bub1-Cdc20-Mad2), likely explaining the brief delay observed in the *ken2* mutant.

### CnBub1 is an active kinase that is required to maintain checkpoint arrests

Even though it has been studied for decades, the importance of Bub1 kinase activity in SAC signalling remains contentious. To test its importance in *Cryptococcus neoformans* we generated a kinase-dead allele (K1011R, D1149N). This was based on a similar kinase-dead mutation made in fission yeast, where it was found that single amino-acid substitutions retained some, albeit very minor Bub1 kinase activity^53^. We have analysed this mutation both *in vivo* and *in vitro*. The full length *bub1-kd* mutant protein was stably expressed in yeast at a comparable level to the wild-type protein (Fig.6a). In Fig.6a the GFP-Bub1 kinase purified from yeast auto-phosphorylates and phosphorylates recombinant histones, in the form of reconstituted nucleosomes, whereas the kinase-dead protein does not. Recombinant Bub1 was purified from *E.coli*. This also phosphorylated itself and phosphorylated H2A on the expected human H2A T120 residue when nucleosomes were used as substrate (Fig. 6b)^53^. These Bub1 kinase results show striking evolutionary conservation: when Bub1 kinase is presented with whole, recombinant nucleosomes, it specifically phosphorylates H2A on the conserved threonine, just as it does in human or fission yeast cells^53^. We expect this phosphorylation of H2A will be relevant to centromere targeting of shugoshin, Aurora B kinase and the CPC, but testing that will be the subject of future CnBub1 studies.

**Figure 6.**
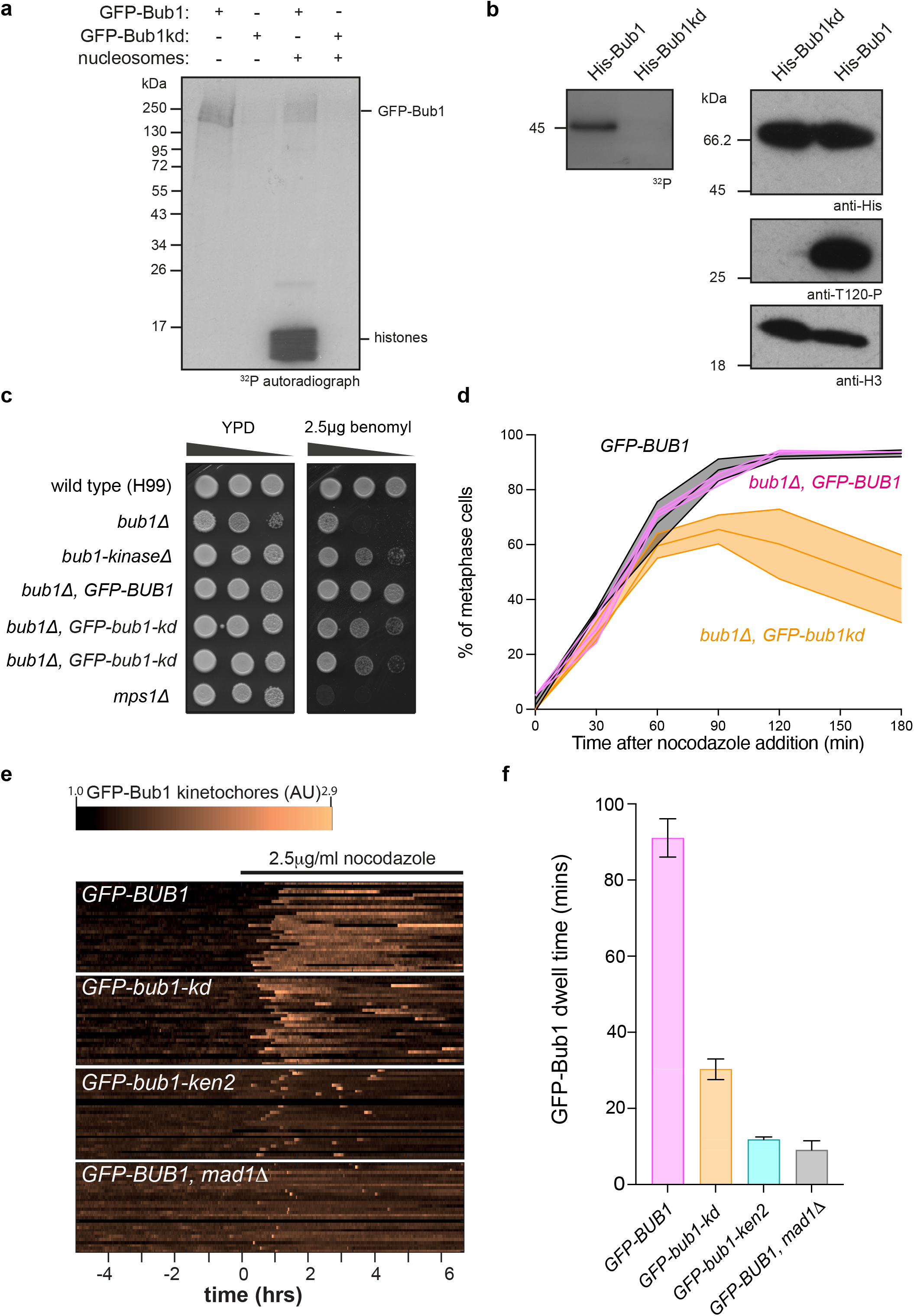
CnBub1 kinase activity is needed to maintain prolonged arrest. a) GFP-Bub1 and *GFP-bub1-kd* were immunoprecipitated from mitotic (nocodazole-arrested) *Cryptococcus* cell extracts using GFP-trap beads, washed and added to kinase assays with or without recombinant nucleosomes as substrate. b) 6xHis-tagged, recombinant Bub1 kinase and a kinase-dead control were purified from bacteria. The kinase-dead protein failed to auto-phosphorylate in the kinase assay (left), nor does it phosphorylate human H2A on Threonine 120 in an immunoblot of a kinase assay with recombinant nucleosomes as substrate (right immunoblots). c) The *bub1* kinase-dead (*-kd*) mutant (see Fig. 6) was expressed in a *bub1Δ* background, and phenotypically compared to the *bub1-kinase truncation* and *mps1* deletion mutants. Strains were diluted, plated and grown on YPGlucose plates with 0 or 2.5μg benomyl (anti-microtubule drug) for 3 days at 30°C. d) Fixed time points were analysed (30, 60, 90, 120 and 180 mins after 2.5μg nocodazole addition) for bright GFP-Bub1 foci (SAC arrested cells). The *bub1-kd* cells delay, but can’t maintain the mitotic arrest. e) Microfluidic temporal heat maps of the indicated strains: nocodazole was added after 5 hours, and cells imaged every 2 mins. f) Plot of the average times with GFP-Bub1 on kinetochores in the SAC delay in these mutant backgrounds (n of 716 for *BUB1-GFP*, 201 for *GFP-bub1-kd*, 109 for *GFP-bub1-ken2*; 199 for *BUB1-GFP, mad1Δ*).

We confirmed that the *bub1-kd* allele was not as sensitive to benomyl as the checkpoint mutant (*mps1Δ*) and that it showed very similar sensitivity to the previously described *bub1* kinase truncated strain^27^ (Fig.6c). We analysed these cells in liquid media to see if they could recruit GFP-bub1-kd to kinetochores and mount a checkpoint response to antimicrotubule drug treatment. Fig.6d analyses cells from fixed time-points (every 30 mins after nocodazole inhibition) for GFP-Bub1 foci, and Fig.6e shows the temporal heat map of kinetochore localization for this mutant from a microfluidic experiment, where we have compared it to the *bub1-ken2* mutant which showed a short mitotic delay. *bub1-kd* cells delayed for longer than the *ken2* allele, for ~30 mins, and then re-budded (see SuppFig3 and Movies). Our interpretation is that Bub1 kinase needs to phosphorylate something to maintain prolonged SAC arrest. There are several possibilities for the relevant SAC substrate, including Cdc20^54,55^. Preliminary *in vitro* kinase assays suggest that recombinant CnCdc20 is a CnBub1 substrate (data not shown). Future work will map these sites in Cdc20 and test whether their modification is important for stabilisation of MCC-APC/C complexes in *Cryptococcus*.

### CnBub1 co-purifies with kinetochores and Cdc20-APC/C

The above data on a series of CnBub1 mutant alleles suggests that CnBub1 likely performs all the functions expected of both Mad3/BubR1 and Bub1 kinase. To test this possibility biochemically we performed large scale purification of GFP-Bub1 from *Cryptococcus* cells, using GFP-Trap magnetic beads, followed by mass spectrometry. We aimed to see whether kinetochore proteins and/or Cdc20-APC/C sub-units could be identified as stable CnBub1 interacting partners.

When we compared the mitotic GFP-Bub1 pull down with an untagged strain (the dashed line represents a significance threshold of P=0.05), we identified kinetochore proteins (Spc105, Stu1), SAC components (Bub3, Mad1, Mad2), SAC effectors (Cdc20, Apc1,2,3,4,5,6,8,10&11) and candidate regulators (PP1, PPZ1) as Bub1-interactors (Fig.7a). We further compared the GFP-Bub1 pull down data from cycling and nocodazole-arrested cell extracts (Fig.7b). Most of the specific interactors associate more strongly with GFP-Bub1 in mitosis. Supporting this, we could co-immunoprecipitate mCherry-Bub1 with GFP-Apc4 from mitotically-arrested *Cryptococcus* extracts (Fig.7c and SuppFig4b).

**Figure 7.**
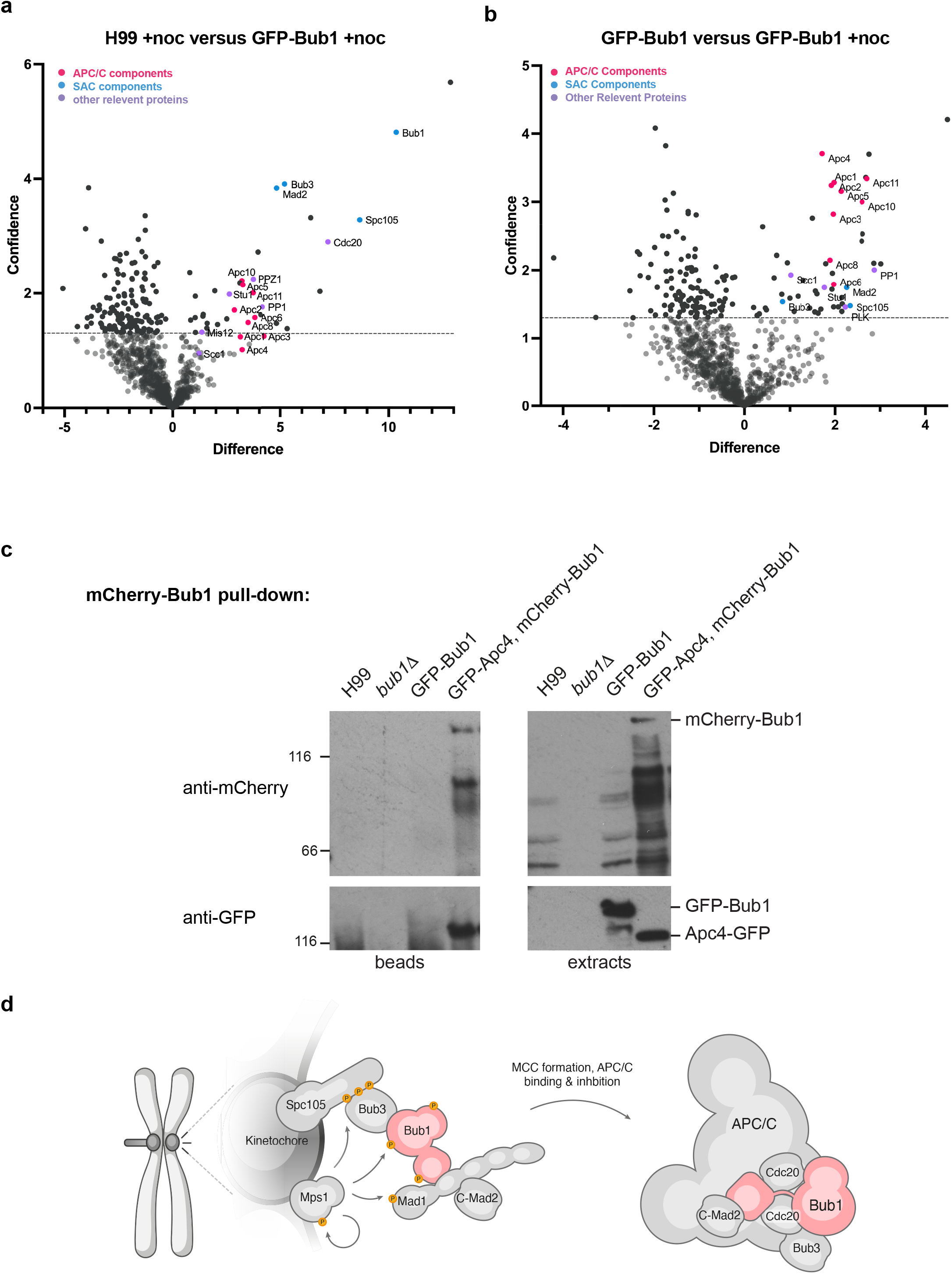
CnBub1 pulls down kinetochores and Cdc20-APC/C. a) GFP-trap immunoprecipitates from a strain expressing GFP-Bub1 were compared to an untagged control strain (H99, wild-type). Both were arrested in mitosis with 2.5μg nocodazole. Immunoprecipitates were run into an SDS-PAGE gel, cut out and digested into peptides with trypsin before analysis on a Orbitrap Fusion™ Lumos™ Tribrid™ Mass Spectrometer. Volcano plots show the difference (mean LFQ difference) and confidence (-log_10_P-value of Perseus statistical test) between the GFP-Bub1 and untagged control pull-downs (n=3 for each). b) Volcano plot showing difference (mean LFQ difference) and confidence (-log_10_(P-value of Perseus statistical test) of GFP-TRAP pull downs from nocodazole-arrested versus cycling GFP-Bub1 cells (n=3 for each). c) Co-immunoprecipitation of GFP-Apc4 (CNAG_04231) with mCherry-Bub1. Strains expressing tagged proteins were arrested in nocodazole and mitotic extracts prepared. mCherry-Bub1 was immunoprecipiated, run on an SDS-PAGE gel and then immunoblotted with anti-GFP and anti-mCherry antibodies, to detect tagged Apc4 and Bub1 respectively. d) Schematic model of the whole SAC pathway summarising the many Bub1 interactions.

Fig.7d shows a schematic model of SAC signalling in *Cryptococcus neoformans* with many of these Bub1 interactions displayed. Our interpretation of this proteomics data is that CnBub1 is recruited to mitotic kinetochores through interaction with phosphorylated Spc105, just as one would expect a Bub1 SAC scaffold. However, such scaffolds are relatively stably bound to the kinetochore, whereas MCC proteins are much more dynamic and exchanging with a soluble pool every few seconds ^56,57^. Preliminary FRAP experiments suggest that the bulk of CnBub1 displays dynamic association with unattached kinetochores (data not shown).

## Discussion

This study has demonstrated striking functional conservation in SAC signalling, even where in *C. neoformans* a single Bub1 polypeptide performs the roles of two proteins in most organisms, Bub1 and Mad3/BubR1. We have demonstrated that CnBub1 is recruited to unattached mitotic kinetochores in a Bub3-mediated fashion, most likely through binding to the kinetochore protein Spc105 after it is phosphorylated by Mps1 kinase. All of this is consistent with CnBub1 having a kinetochore-based signalling scaffold role. Yet proteomics, structural modelling and site-directed mutagenesis all argue that the CnBub1 protein also has the key downstream effector function, interacting with Cdc20-APC/C to delay the onset of anaphase. Thus, we have confirmed that CnBub1 performs both Bub1 and Mad3/BubR1 SAC functions. Our findings are consistent with the observation that the single *L.kluyveri* MadBub gene can rescue a *S.cerevisae bub1,mad3* double deletion strain^58^.

To analyse the importance of distinct mechanistic functions in more detail, we have developed a novel micro-fluidics SAC assay. This enables imaging of single cells in a range of media and conditions, and measures the time spent in each mitotic cycle with GFP-Bub1 at kinetochores. Upon nocodazole treatment, this was very revealing and enabled quantitative analysis of a range of mitotic delays in different *bub1* alleles. The *bub1-cd1, ken1* and *ken2* phenotypes are remarkably consistent with what one might predict from studies of model systems, such as *S.pombe, S.cerevisiae* and human cells over the past 30 years. Very little (if any) SAC delay remains when either Cdc20 (MCC) or Mad1 binding are disrupted, through mutation of KEN1 or CD1. After mutation of KEN2, where Cdc20-APC/C can’t be stably bound by CnBub1, a mitotic delay of ~10 mins was still observed (see Fig.5b and 6f). Removing the kinase activity of Bub1 has the weakest phenotype and a SAC delay of ~30 mins was still apparent. This finding is consistent with studies in other organisms (and our own unpublished work in *S. pombe*) where Bub1 kinase activity is only required to maintain prolonged SAC arrest, most likely through phosphorylation of Cdc20^54,55^. More generally, related micro-fluidics assays will be an excellent way to study single cell responses of *Cryptococcus neoformans* to a whole range of stresses.

We conclude that whilst many aspects of mitosis are distinctly different in *Cryptococcus* (a partially open mitosis, scattered KTs and MTOCs in interphase, and distinct ploidy shifts^23^) the molecular mechanism of SAC signalling is remarkably well conserved. Thus most of what has been learnt from studying the SAC in model systems and humans, is very likely relevant to studies of this important fungal pathogen. It will be fascinating to see how important SAC signalling and other Bub1 functions are in polyploid Titan cell divisions.

Preliminary FRAP studies suggest that the bulk of CnBub1 (~80%) exchanges rapidly on mitotic kinetochores, consistent with behaviour of BubR1 and Cdc20 at human kinetochores^56,57^. Whether all of the CnBub1 is dynamic, or there is also a small pool acting as a stable scaffold at kinetochores will require careful analysis. It will be very interesting to see whether the APC/C also interacts with kinetochores in a dynamic fashion, exchanging with a diffuse nuclear pool.

### Non-SAC functions for Bub3-Bub1?

A striking observation is the severity of the complete loss of function phenotype of CnBub1 and CnBub3 (see Fig.1). These mutants are much sicker than *mad* and *mps1* mutants, indicating that the Bub complex is likely to have important functions outside of core SAC signalling. We have demonstrated that CnBub1 kinase phosphorylates nucleosomes on the conserved T120 residue of human H2A *in vitro*. In other systems this chromatin mark helps recruit Sgo1 and CPC to kinetochores^53,59,60^. We are currently testing the hypothesis that Bub1 kinase carries out related bi-orientation and error-correction roles in *Cryptococcus* mitosis. In addition, CnBub1 could be involved in cross-talk with other stress and checkpoint signalling responses. *Cnbub1* mutants were previously shown to be sensitive to hydrogen peroxide and 5-Flucytosine^27^. Much remains to be learnt about the many roles of CnBub1, but we believe that it is already a relevant anti-mitotic drug target and we are pursuing a search for small molecule inhibitors of this fascinating mitotic protein.

## Acknowledgements

We would like to thank all members of the Hardwick and JP labs for their support, discussions and suggestions on this manuscript; Paige Erpf and James Fraser for Safe Haven and Blaster constructs; Kaustuv Sanyal, Lukasz Kozubowski and Liz Ballou for *Cryptococcus neoformans* strains and plasmids; Ken Sawin for anti-mCherry antibodies; Connie Nichols for information on pCN19; Liz Ballou for many helpful *Cryptococcus* tips and suggestions; Dave Kelly for help with the microscopy and generating scripts and Peter Swain for leading on the ISSF funding (IC).

This work was supported by grants from the Leverhulme Trust (RPG-2018-379 to IL, KGH); the Darwin Trust of Edinburgh (KA, APS); the Wellcome Trust (SRF 202811 to AAJ; WCCB core grant 203149; iCM programme 218470 to TD, and the Wellcome Trust-University of Edinburgh Institutional Strategic Support Fund, IC) and 2021R1A2B5B03086596 from the National Research Foundation of Korea (NRF) funded by the Ministry of Science and ICT (MSIT) (Y-SB).

## Methods

***Cryptococcus* strains** were all derived from *Cryptococcus neoformans* var. *grubii* H99 YSB3632 *mps1Δ::NAT* (*Ref 27*)

YSB3398 *bub1Δ::NAT* (*this study*)

YSB4190 *bub1-kinase domain truncation::NAT* (*Ref 27*)

CNSD159 *TUB4::TUB4-mCherry:G418* (*Ref 24*)

CNVY105 *DAD2::DAD2-mCherry:G418* (*Ref 24*)

KA51 *mad1Δ* (amdS2 Blaster)

IL006 *mps1Δ::NAT, HISp:GFP-Bub1:HYG* (chromosome 3, safe haven p37)

IL089 *mad1Δ, HISp:GFP-Bub1:HYG* (chrom 3, safe haven p37)

IL102 *bub1Δ::NAT, HISp:GFP-Bub1:HYG* (chrom 3, safe haven p37)

IL143 *bub1Δ::NAT, HISp:GFP-bub1-kd:HYG* (chrom 3, safe haven p37)

IL173 *bub1Δ::NAT, HISp:GFP-bub1-cd1:HYG* (chrom 3, safe haven p37)

IL178 *bub1Δ::NAT, HISp:GFP-bub1-glebs:HYG* (chrom 3, safe haven p37)

IL226 *bub1Δ::NAT, HISp:GFP-bub1-ken1:HYG* (chrom 3, safe haven p37)

IL231 *bub1Δ::NAT, HISp:GFP-bub1-ken2:HYG* (chrom 3, safe haven p37)

IL066 *GAL7p:HA-Bub3:HYG* (chrom 3, safe haven p37)

IL111 *bub3Δ, GAL7p:HA-Bub3:HYG* (chrom 3, safe haven p37)

IL187 *Dad2-RFP:G418, HISp:GFP-Bub1:HYG* (chrom 3, safe haven p37)

IL194 *Tub4-RFP:G418, HISp:GFP-Bub1:HYG* (chrom 3, safe haven p37)

IL129 *bub1Δ::NAT, HISp:GFP-Bub1:HYG* (chrom 3, safe haven p37), *GPDIp:mcherry-Bub3:G418* (chrom 14, safe haven p36)

IL86 *HISp:GFP-Bub1:HYG* (chrom 3, safe haven p37), *GPDIp:mcherry-Bub3:G418* (chrom 14, safe heaven p36)

IL305 *bub3Δ, GAL7p:HA-Bub3:HYG* (chrom 3, safe haven p37), HISp:mcherry-Bub1:NATR (chrom 7, safe heaven 31)

IL278 *bub1Δ::NAT, HISp:mcherry-Bub1:HYG* (chrom 3, safe haven p37), *HISp:GFP-APC4* (chrom 7, safe haven 37).

### Yeast media

All *Cryptococcus* strains were generated in the H99 genetic background and stored in YPDA (2% bactopeptone, 1% yeast extract, 2% glucose, 2% agar, 1mM adenine) with 50% glycerol at −80 °C. Strains were grown on YPDA plates for two days prior to use. For microfluidic experiments yeast synthetic complete media (SC: 0.67% Yeast Nitrogen Base; amino acid mix; 2% glucose). Knockout transformants generated using the blaster cassette were selected on plates containing YNB medium (0.45% yeast nitrogen base w/o amino acids and ammonium sulphate, 2% glucose, 10mM ammonium sulphate, 2% agar, 1mM adenine) with 5mM acetamide. To get rid of the blaster cassette strains were restreaked on YNB media containing 2% glucose, 10mM ammonium sulphate, and 10mM fluoroacetamide, as described previously^30^. For drug selection cells were grown on YPDA plates containing 300μg/ml hygromycin B (Invitrogen), 100μg/ml Geneticin (G418 Sulphate, Gibco) or 100μg/ml nourseothricin (Jena Bioscience). For expression from the *GAL7* promoter: this promoter is switched on and off by adding 2% galactose or 2% glucose respectively.

### Gibson assembly, sequencing

Primers were purchased from Sigma/Merck or IDT. Gibson assembly and NEB builder assembly were performed following manufacturer’s instructions; sequencing was carried out using Big Dye.

#### Plasmids generated in this study

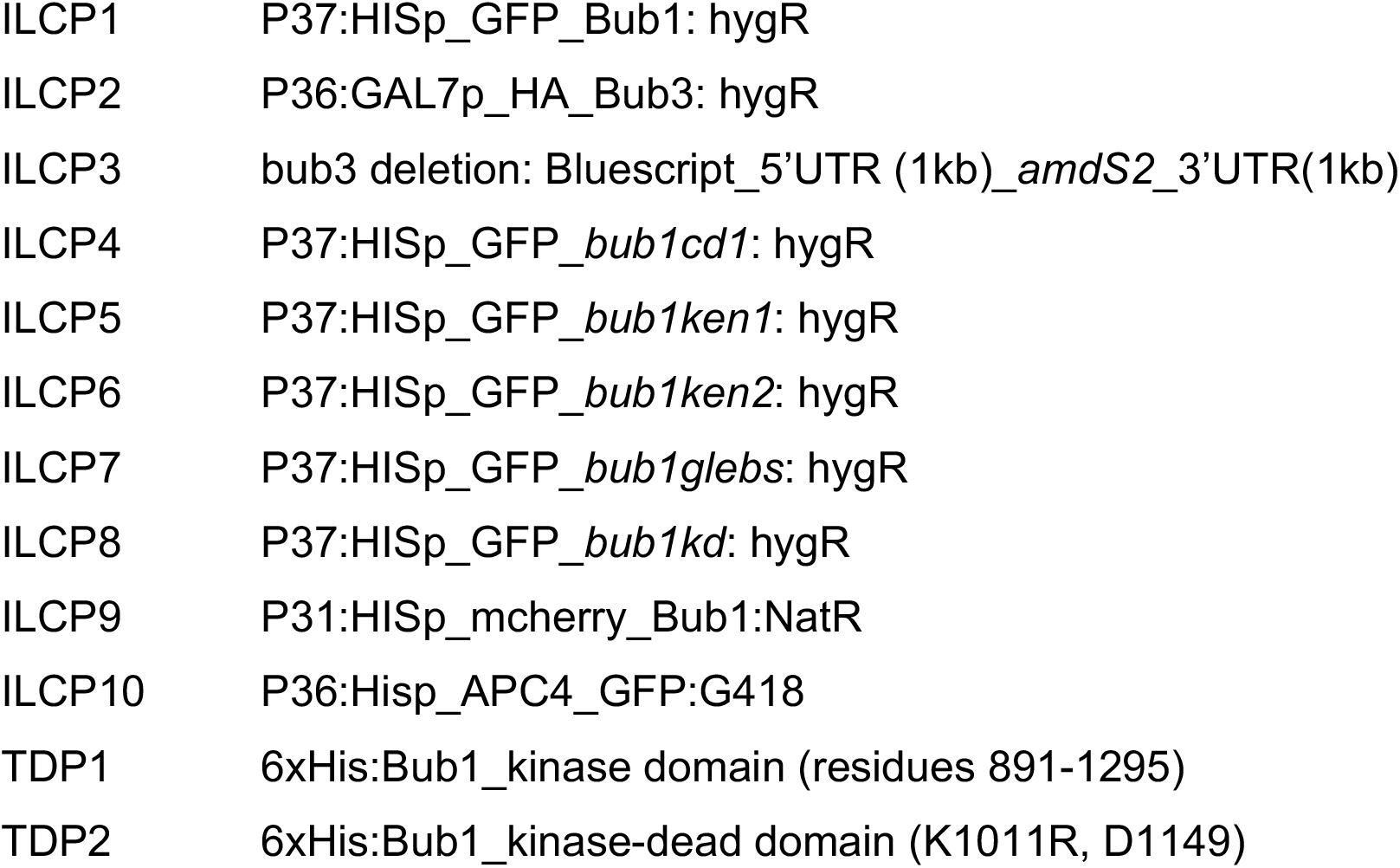

**Plasmids generated in this study:**
|  |  |
| --- | --- |
| ILCP1 | P37:HISp_GFP_Bub1: hygR |
| ILCP2 | P36:GAL7p_HA_Bub3: hygR |
| ILCP3 | bub3 deletion: Bluescript_5'UTR (1kb)_amdS2_3'UTR(1kb) |
| ILCP4 | P37:HISp_GFP_bub1cd1: hygR |
| ILCP5 | P37:HISp_GFP_bub1ken1: hygR |
| ILCP6 | P37:HISp_GFP_bub1ken2: hygR |
| ILCP7 | P37:HISp_GFP_bub1glebs: hygR |
| ILCP8 | P37:HISp_GFP_bub1kd: hygR |
| ILCP9 | P31:HISp_mcherry_Bub1:NatR |
| ILCP10 | P36:Hisp_APC4_GFP:G418 |
| TDP1 | 6xHis:Bub1_kinase domain (residues 891-1295) |
| TDP2 | 6xHis:Bub1_kinase-dead domain (K1011R, D1149) |

#### Primers used in this study

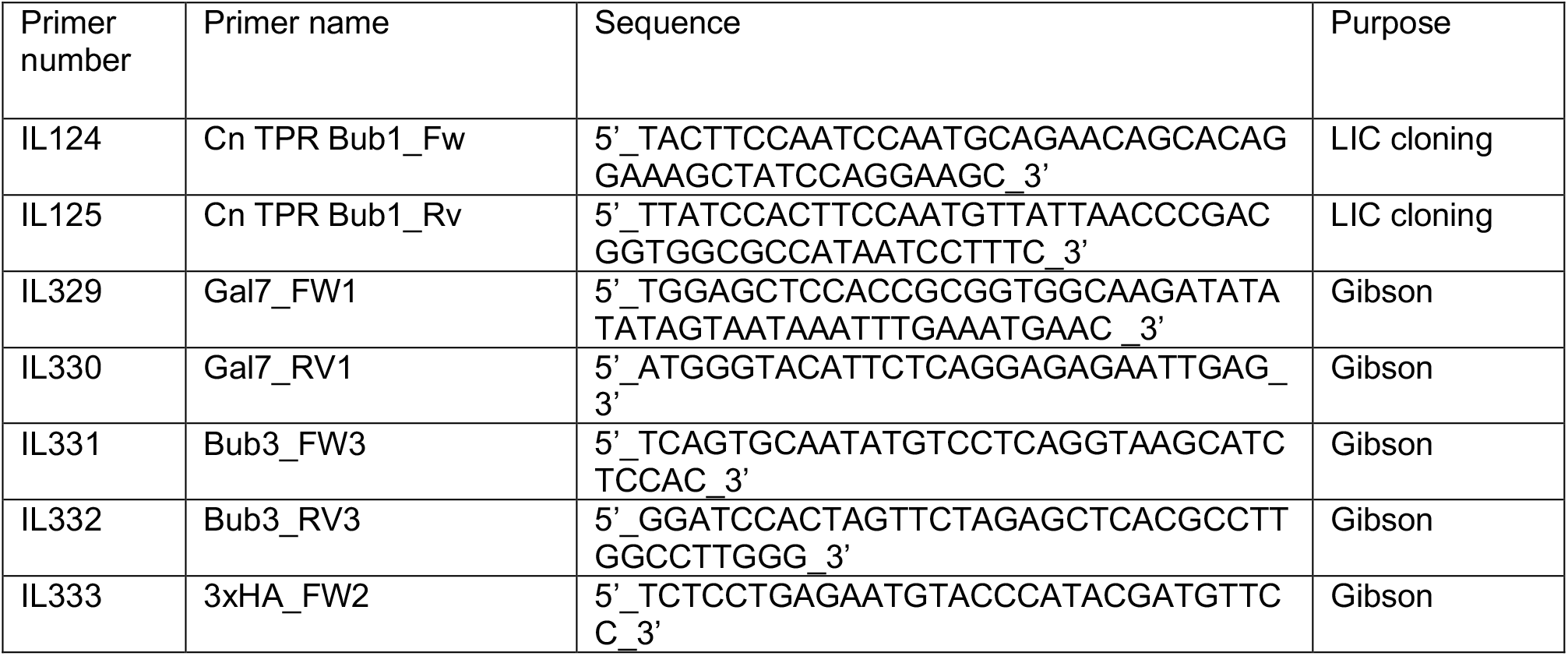
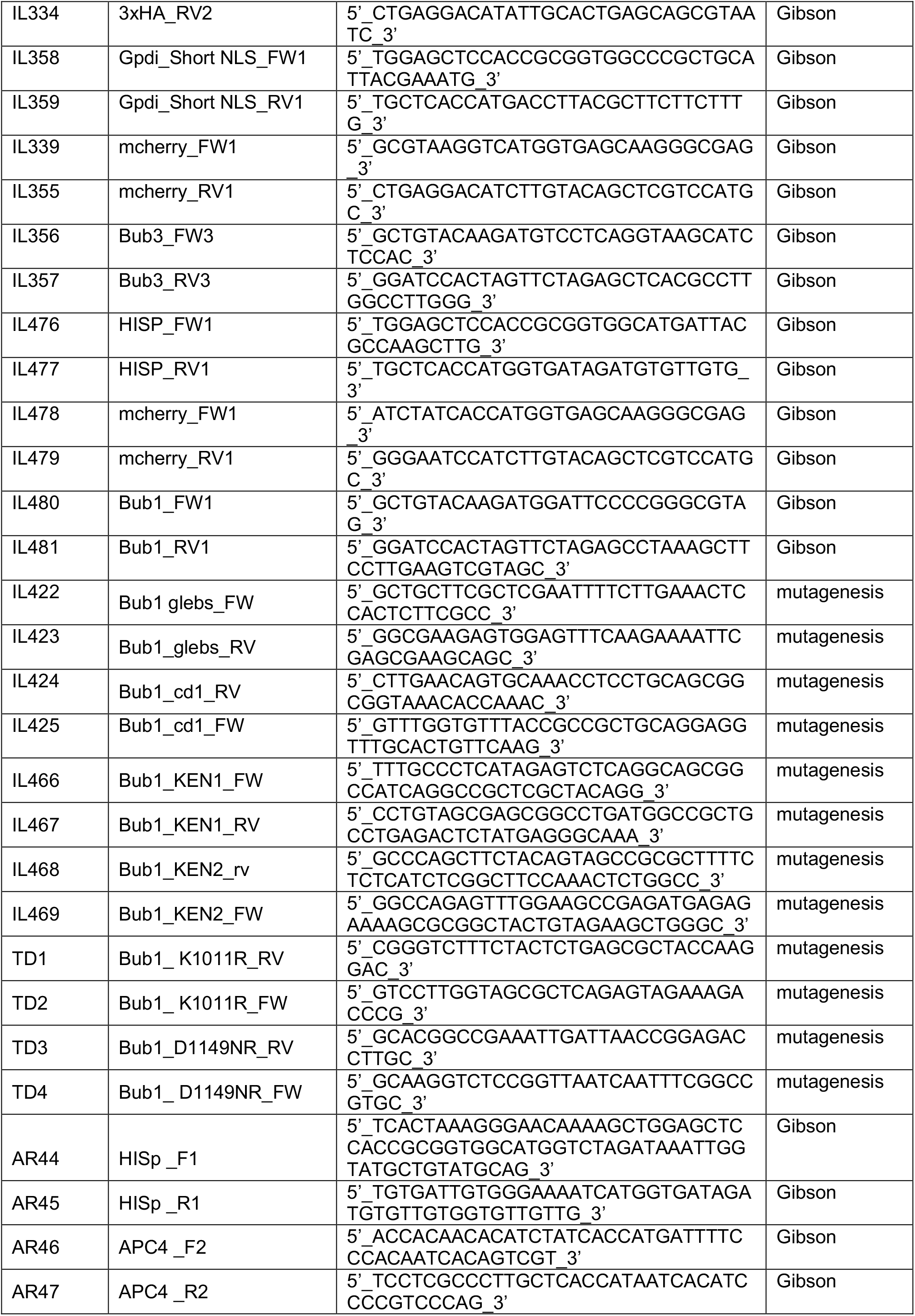
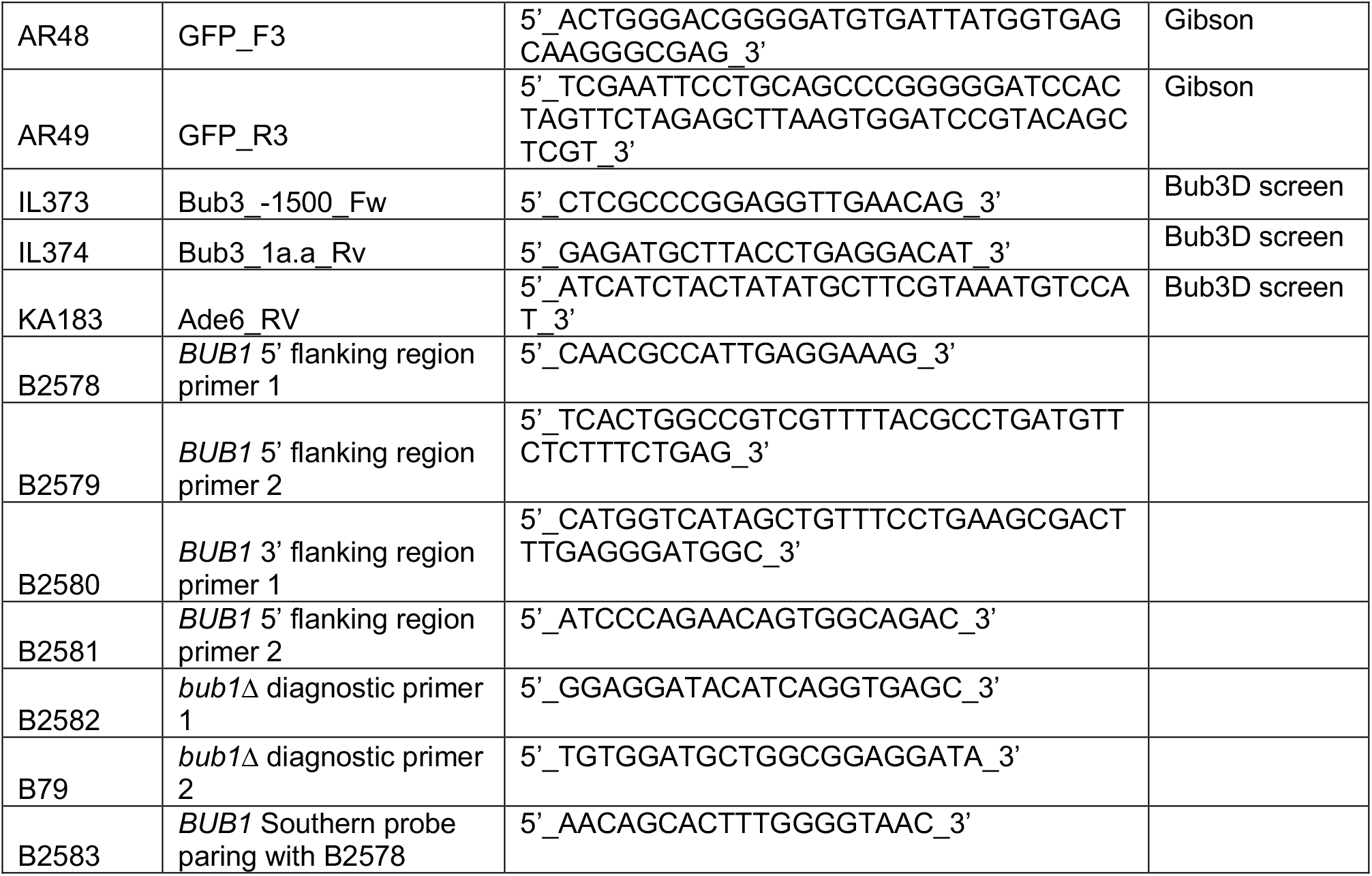

### Bub1 knockout and rescue (HISp-GFP-Bub1)

The complete Bub1 knockout was made homologous recombination of the PCR amplified *BUB1::NAT* knockout cassette. Briefly, the amplified 5’ and 3’ flanking regions were fused to split 5’ and 3’ nourseothricin-resistance markers (*NAT*; nourseothricin acetyl transferase), respectively, by double-joint PCR as previously reported^24^. The knockout cassette was introduced into H99 wild-type strain by biolistic transformation and selected on YPD medium containing 100μg/mL nourseothricin. The transformants were confirmed by diagnostic PCR and Southern blot analysis (SuppFig 1a).

### HISp-GFP-Bub1

For construction of HISp-GFP-Bub1 the pCN19 vector was digested with SpeI and ligated with a full length Bub1 clone that had been PCR-amplified from H99 genomic DNA. The resulting construct was digested with XbaI and NruI and the p37 safe-haven vector (developed by James Fraser^30,39^) digested with Xbal and EcoRV, then the linearised fragments ligated. Plasmids were sequenced and the final vector digested with PacI to target homologous recombination to the correct chromosomal safe-haven locus.

### Bub1 alleles (kinase-dead, KEN1&2, GLEBS, CD1)

The HISp-GFP-Bub1 plasmid was mutagenised using the Quickchange lightning kit (Agilent), according to the manufacturer’s instructions.

### GPDIp-mCherry-Bub3

The GPDI promoter, mCherry and Bub3 genomic fragments were assembled into p36 (G418 resistance, chromosome 14 safe-haven) using Gibson assembly.

### *Blaster knockout of BUB3 and the* P_GAL_-*BUB3 ‘rescue’ construct*

Ectopic expression of HA-tagged Bub3 was generated under the P_GAL7_ promoter ^41^.^61^. The Gal7 promoter, the HA tag and the Bub3 genomic fragment were inserted into pPEE37 (James Fraser) using Gibson assembly (NEB). Expression of HA-Bub3 was confirmed by immunoblotting with anti-HA antibodies. Due to the very strong phenotype of *bub3Δ* the endogenous gene was deleted in this strain, already containing the ‘rescue construct’, GAL-HA-Bub3. The Bub3 knockout construct was made using the Blaster construct ^30^. 1 kb homologous arms, consisting of the 5’ and 3’ Bub3 UTR sequences, were ligated at both ends of the selective *amdS2* marker: BlueScript vector was digested with HindIII/EcoRI, and pEE8 with SpeI/HindIII. The final construct was digested with SacII and XhoI, transformed into IL66, and colonies selected on acetamide media. Cells were restreaked several times to ensure stable integration had occurred and that all cells contained the Blaster. Correct integration of the marker was confirmed by PCR analysis of genomic DNA made from stable transformants (SuppFig.1b).

### Cn transformation protocol

Strains were pre-cultured in YPDA, then diluted overnight into 100ml of culture. Overnight cultures were grown at 30°C (25 °C for *bub1Δ*) until the OD_600_ reached 0.3-0.36 the next morning. Cells were harvested, washed twice with 50ml iced cold water and once with 50ml of ice-cold electroporation buffer [10mM Tris-HCl pH7.5, 1mM MgCl_2_ and 270mM sucrose]. Then cells were resuspended in 35ml electroporation buffer containing 150μl of 1M DTT and incubated for 15 mins on ice. Cells were then harvested again, washed with 50ml electroporation buffer and finally resuspended in ~100μl of electroporation buffer. 40μl of cell suspension was gently mixed with linearised DNA (~4μg) and transferred to pre-cooled electroporation cuvettes (2mm). Electroporation was performed at 1400 V, 600 ohm and 25μF (Biorad, GenePulser). Following electro-pulse cells were incubated on ice for two minutes then 1 ml pre-warmed (30 °C) YPDA was added to the cells before plating on YPDA. After overnight recovery, cells were replica-plated on selection plates. Colonies were typically tested by PCR for correct construct integration and by western blot for protein expression.

### Bub1 antibody generation and purification

Residues 53-209 of Bub1 were amplified from cDNA and cloned into a Biobrick vector (9B, pET His6 TEV, a gift from Scott Gradia, https://www.addgene.org/48284/). This Bub1 construct was expressed in *E. coli* ArcticExpress cells (Agilent), purified on Cobalt resin, eluted and then dialysed into 50mM Hepes pH7.6, 75mM KCl. This recombinant protein was used to immunise sheep (MRC PPU Reagents and Services, University of Dundee). Specific antibodies were affinity-purified using Affigel 10 resin coupled to the same recombinant 6xHis-Bub1 protein. Sheep sera was diluted with PBS, filtered, then gently pumped through the Affigel-Bub1 column overnight. The column was then thoroughly washed with PBS-Tween, then PBS containing an additional 0.5M NaCl, and finally eluted with 100 mM triethylamine (pH 11.5), before dialysing the antibodies overnight into PBS containing 40% glycerol.

### P_HIS_-GFP-Bub1 and mCherry-Bub3 microscopy

Live-cell microscopy was performed with a Spinning Disc Confocal microscope (Nikon Ti2 CSU-W1) with a 100X oil objective (Plan Apo VC) coupled to a Teledyne-Photometrics 95B sCMOS camera. For imaging, Z-stacks of 11 images (step size 0.5μm) were acquired using a 491nm laser line for GFP and 561nm laser for m-Cherry. Exposure times were 300ms and laser power was kept to the minimum to avoid photobleaching. Images were captured using Nikon Elements software.

ImageJ was used for all image analysis and specific scripts for GFP-Bub1 cell counting can be found here (web-link to be added). Images were further processed in Adobe Photoshop to adjust brightness and contrast, all adjustments were applied to whole images uniformly, and to all images being compared.

### Checkpoint assays

#### Benomyl plates - serial dilution assay

Cells from an overnight culture were diluted to OD_600_ ~0.4 in distilled water. 10-fold, serial dilutions were made and spotted onto YPDA plates (with or without the anti-microtubule drug, benomyl at different concentrations) then typically incubated at 30°C for 48 hours. Benomyl stock was 30mg/ml in DMSO, and due to solubility issues this was added directly to boiling YPD agar.

### Time lapse microfluidic assays

The utility of microfluidics for single cell analysis in *Cryptococcus* was demonstrated previously in studies of ageing^62^. We used the Alcatras cell traps^49^ incorporated into devices allowing for use with multiple strains^63,64^. We moulded devices in polydimethylsiloxane (PDMS) from an SU8-patterned wafer with an increased thickness of 7μm, to accommodate the larger size of *C neoformans* cells compared to *S cerevisiae* (manufactured by Microresist, Berlin, design available on request). Imaging chambers for individual strains are isolated by arrays of PDMS pillars separated by 2μm gaps. This prevents intermixing of strains while cells experience identical media conditions.

Before use we filled the devices with synthetic complete (SC) media, supplemented with 0.2g/l glucose and containing 0.05%w/v bovine serum albumin (Sigma) to reduce cell-cell and cell-PDMS adhesion. Cells pre-grown to logarithmic phase in the same media (lacking the BSA) were injected into the device. An EZ flow system (Fluigent) delivered media at 10μl per minute to the flow chambers and performed the switch to media containing nocodazole after 5 hours. This media also contained Cy5 dye to allow monitoring of the timing of the media switch. We captured image stacks at 2_minute intervals at 4 stage positions for each strain, using a Nikon TiE epifluorescence microscope with a 60x oil-immersion objective (NA 1.4), a Prime95b sCMOS camera (Teledyne Photometrics) and OptoLED illumination (Cairn Research). Image stacks had 5 Z-sections, separated by 0.6μm, captured using a piezo lens positioning motor (Pi).

### Time lapse image processing and analysis

Cell outlines were segmented using a method developed by the Swain laboratory based on a convolutional neural network (manuscript in preparation, details available on request). Matlab (Mathworks) was used for further processing and visualisation. To quantify kinetochore dwell time of Bub1-GFP fluorescence we create a projection of the maximum values from all GFP sections then divide the median fluorescence of the brightest 5 pixels within each cell by the median fluorescence of the cell as a whole. This ratio gives a reliable measure of the aggregation of protein at an organelle and has been used extensively for quantifying nuclear localization^65–67^. In the absence of nocodazole, cells that have any outlier values for the ratio, defined as values greater than 3 scaled median absolute deviations away from the median, are removed. Cells present for fewer than 80% of the time points recorded are also removed.

For each strain, the ratios were normalised by dividing by the mean value for wild type cells in the absence of nocodazole, measured in the same experiment. For calculation of the kinetochore dwell times (Fig.5b and 6f), a threshold of 1.22 was applied to the ratio values at each time point to determine the presence or absence of Bub1 at the kinetochore. Small gaps in localization (due to kinetochore signal dropping briefly out of focus) were removed by applying a 1-dimensional morphological closing to the thresholded image, with a 3 pixel (3 timepoint or 6 minutes) structuring element. For the temporal heat maps (Fig.5a and 6e) the data for 30 cells present for the whole experiment are selected for inclusion from each strain pseudorandomly using the Matlab rand function.

#### Immunoblot analysis

For whole-cell lysates, typically a 10 ml (OD_600_ ~0.5) cell culture was harvested by centrifigation. Cells were washed once and snap frozen in liquid nitrogen. Frozen pellets were resuspended in 2x sample buffer with 200mM DTT and 1mM PMSF. Cells were then disrupted with 0.5mm Zirconia/Silica beads (Thistle Scientific) using a multi-bead beater for 1 min (BioSpec Products). Samples were spun, boiled for 5 min at 95°C, and the cell debris pelleted by centrifugation for 5 min at 13000 rpm. Cleared extracts were then immediately loaded and separated by SDS-PAGE. Proteins were transferred onto nitrocellulose membrane (Amersham Protran 0.2 μm nitrocellulose, GE Healthcare Lifescience) using a semi-dry transfer unit (TE77, Hoefer, Inc, MA, USA) in 25mM Tris, 130 mM glycine and 20% methanol. Transfer was typically for 1.5-2 hours at 150-220 mA. Efficiency of protein transfer was visualised using Ponceau S solution. Membranes were blocked with Blotto (PBS, 0.04% Tween 20, 4% Marvel skimmed milk powder) for at least 30 min at room temperature on a shaking platform. The primary antibody (anti-Bub1, anti-GFP or anti-HA) was incubated in the same blocking buffer (1 in 1000 dilution) overnight at 4°C. The membrane was then washed 3 times for 10 mins with PBS+0.04% Tween and then incubated with the secondary antibody (1 in 5000 dilution) for at least an hour at room temperature. The membrane was washed again, rinsed with PBS and ECL performed (SuperSignal West Pico, or SuperSignal West Femto, Thermo Fisher Scientific Inc, IL, USA).

### Bub1 purifications and kinase assays

Residues 891-1295 of CnBub1 were amplified from cDNA and cloned into the pET His6 TEV 9B BioBrick cloning vector. Induction of protein expression was performed in BL21 (pLysS) cells. IPTG was added and cultures incubated for 16 hrs at 18°C. Cells were harvested, washed and pellets frozen in liquid nitrogen. Cell pellets were resuspended in lysis buffer [50mM Tris-HCl pH8, 500mM NaCl, 10% Glycerol, 5mM Imidazole, 1mM β-mercaptoethanol, EDTA-free protease inhibitor tablet (Roche), 1mM PMSF] then lysed by sonication (1 sec ON and 2sec OFF for a total of 3 min). To remove the cell debris, lysed cells were centrifuged at 20,000 rpm, for 30-45 min, at 4°C, and the lysate filtered through a 0.45μm syringe. Lysates were then incubated with rotation for 2 hours (at 4°C) with Talon cobalt resin (Thermofisher). After incubation, the beads were transferred to a Biorad column, washed with 10 column volumes of wash buffer, and protein eluted with lysis buffer containing 250mM imidazole. The recombinant kinase domain was dialysed overnight into 50mM Tris-HCl pH8, 150mM NaCl, 5% glycerol, 2mM DTT.

Protein was concentrated via centrifugation (Vivaspin) and activity assayed against reconstitutes nucleosome substrates for phosphorylation of T120 residue of Histone H2A. Recombinant nucleosomes were purified as described previously ^68^. Recombinant kinase was added to 10μl of 2X kinase buffer [40mM Hepes (pH 7.5), 200mM KCl, 20mM MgCl_2_, 2mM DTT, 400μM ATP] and nucleosomes, and water to a final volume of 20μl. Reactions were incubated at 30°C for 30 min and quenched with an equal volume of SDS-PAGE sample buffer, and run on an SDS-PAGE gel. Immunoblot analysis was performed as above, with anti-His and anti-T120 phosphoantibody (Active motif, 39391).

Radioactive Bub1 kinase assays were performed in a similar reaction for 30 min at 30°C: 20mM Hepes (pH 7.5), 100mM KCl, 10mM MgCl_2_, 1mM DTT, 100μM cold ATP, 5μCi γ-^32^P-labelled-ATP and 1μg recombinant nucleosomes.

#### Lysis of large scale cell extracts for mass spectrometry

Yeast cells were grown overnight (to OD_600_ of ~0.5) in 500mls of YPDA. 2.5μg/ml nocodazole was added to the cells and incubated for three hours. Cells were harvested by centrifugation at 5000 rpm at 4°C, for 15 mins. Pelleted cells were frozen in drops, using liquid nitrogen. The cells were then ground manually, using a mortar and pestle. Yeast powders were resuspended into lysis buffer containing 50mM Hepes pH7.6, 75mM KCl, 1mM MgCl_2_, 1mM EGTA, 10% Glycerol, 0.1% Triton X-100, 1mM Na3VO4, 10μg/mL CLAAPE (protease inhibitor mix containing chymostatin, leupeptin, aprotinin, antipain, pepstatin, E-64 all dissolved in DMSO at a final concentration of 10 mg/mL), 1 mM PMSF, 0.01 mM microcystin. 1g of yeast powder was resuspended in 1ml of the lysis buffer. Cell lysis was completed by sonication (cycles of 5 sec ON, 5 sec OFF for 1 min). After sonication, the cell debris was pelleted (30 min, at 22000 rpm, at 4°C) and the supernatant incubated with anti-GFP TRAP magnetic agarose beads (ChromoTek) for 1 hr at 4°C. The beads were washed at least 9 times with wash buffer (50mM Hepes pH7.6, 75mM KCl, 1mM MgCl_2_, 1mM EGTA, 10% Glycerol) and once with PBS+0.001% Tween 20. Proteins were eluted from the beads by adding 2X sample buffer containing 200mM DTT and boiled at 95°C for 5-10 min, before running on an SDS-PAGE gel.

### GFP-Bub1 mass-spectrometry and volcano plots

Protein samples from all biological replicates were processed at the same time and using the same digestion protocol without any deviations. They were subjected for MS analysis under the same conditions, and protein and peptide lists were generated using the same software and the same parameters. Specifically, proteins were separated on gel (NuPAGE Novex 4-12% Bis-Tris gel, Life Technologies, UK), in NuPAGE buffer (MES) and visualised using Instant*Blue*™ stain (AbCam, UK). The stained gel bands were excised and de-stained with 50mM ammonium bicarbonate (Sigma Aldrich, UK) and 100% (v/v) acetonitrile (Sigma Aldrich, UK) and proteins were digested with trypsin, as previously described ^69^. In brief, proteins were reduced in 10mM dithiothreitol (Sigma Aldrich, UK) for 30mins at 37°C and alkylated in 55mM iodoacetamide (Sigma Aldrich, UK) for 20 min at ambient temperature in the dark. They were then digested overnight at 37°C with 12.5 ng trypsin per μL (Pierce, UK). Following digestion, samples were diluted with an equal volume of 0.1% TFA and spun onto StageTips as described previously ^70^. Peptides were eluted in 40 μL of 80% acetonitrile in 0.1% TFA and concentrated down to 1 μL by vacuum centrifugation (Concentrator 5301, Eppendorf, UK). The peptide sample was then prepared for LC-MS/MS analysis by diluting it to 5 μL with 0.1% TFA.

LC-MS analyses were performed on an Orbitrap Fusion™ Lumos™ Tribrid™ Mass Spectrometer (Thermo Fisher Scientific, UK) coupled on-line, to an Ultimate 3000 HPLC (Dionex, Thermo Fisher Scientific, UK). Peptides were separated on a 50 cm (2 μm particle size) EASY-Spray column (Thermo Scientific, UK), which was assembled on an EASY-Spray source (Thermo Scientific, UK) and operated constantly at 50°C. Mobile phase A consisted of 0.1% formic acid in LC-MS grade water and mobile phase B consisted of 80% acetonitrile and 0.1% formic acid. Peptides were loaded onto the column at a flow rate of 0.3 μL min^−1^ and eluted at a flow rate of 0.25 μL min^−1^ according to the following gradient: 2 to 40% mobile phase B in 150 min and then to 95% in 11 min. Mobile phase B was retained at 95% for 5 min and returned back to 2% a minute after until the end of the run (190 min). Survey scans were recorded at 120,000 resolution (scan range 350-1500 m/z) with an ion target of 4.0e5, and injection time of 50ms. MS2 was performed in the ion trap at a rapid scan mode, with ion target of 2.0E4 and HCD fragmentation (Olsen *et al*, 2007) with normalized collision energy of 28. The isolation window in the quadrupole was 1.4 Thomson. Only ions with charge between 2 and 6 were selected for MS2. Dynamic exclusion was set at 60 s.

The MaxQuant software platform ^71^ version 1.6.1.0 was used to process the raw files and search was conducted against our in-house *Cryptococcus neoformans* var. *grubii* protein database, using the Andromeda search engine ^72^. For the first search, peptide tolerance was set to 20 ppm while for the main search peptide tolerance was set to 4.5 pm. Isotope mass tolerance was 2 ppm and maximum charge to 7. Digestion mode was set to specific with trypsin allowing maximum of two missed cleavages. Carbamidomethylation of cysteine was set as fixed modification. Oxidation of methionine, and phosphorylation of serine, threonine and tyrosine were set as variable modifications. Label-free quantitation analysis was performed by employing the MaxLFQ algorithm ^73^. Peptide and protein identifications were filtered to 1% FDR. Statistical analysis was performed by Perseus software ^74^, version 1.6.2.1.

## MS data availability

(links to PRIDE will be added here)

## Supplementary data files (Leontiou et al)

**Supp Figure 1.**
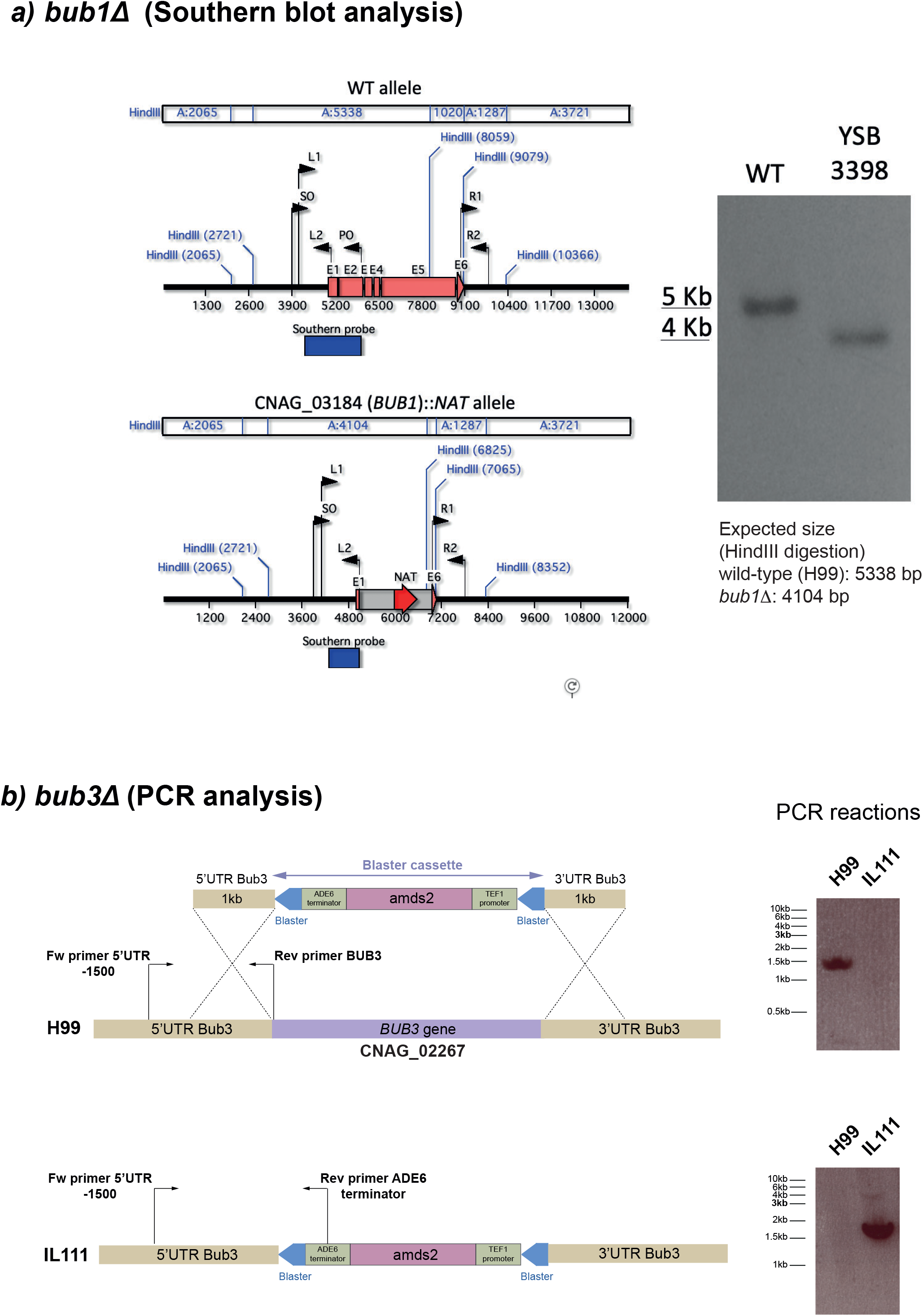
a) Schematic of *bub1* deletion construction and Southern blot. b) Schematic of *bub3* deletion construction using the amds2 Blaster cassette, indicating the primers used to confirm replacement of the *BUB3* gene.

**Supp Figure 2.**
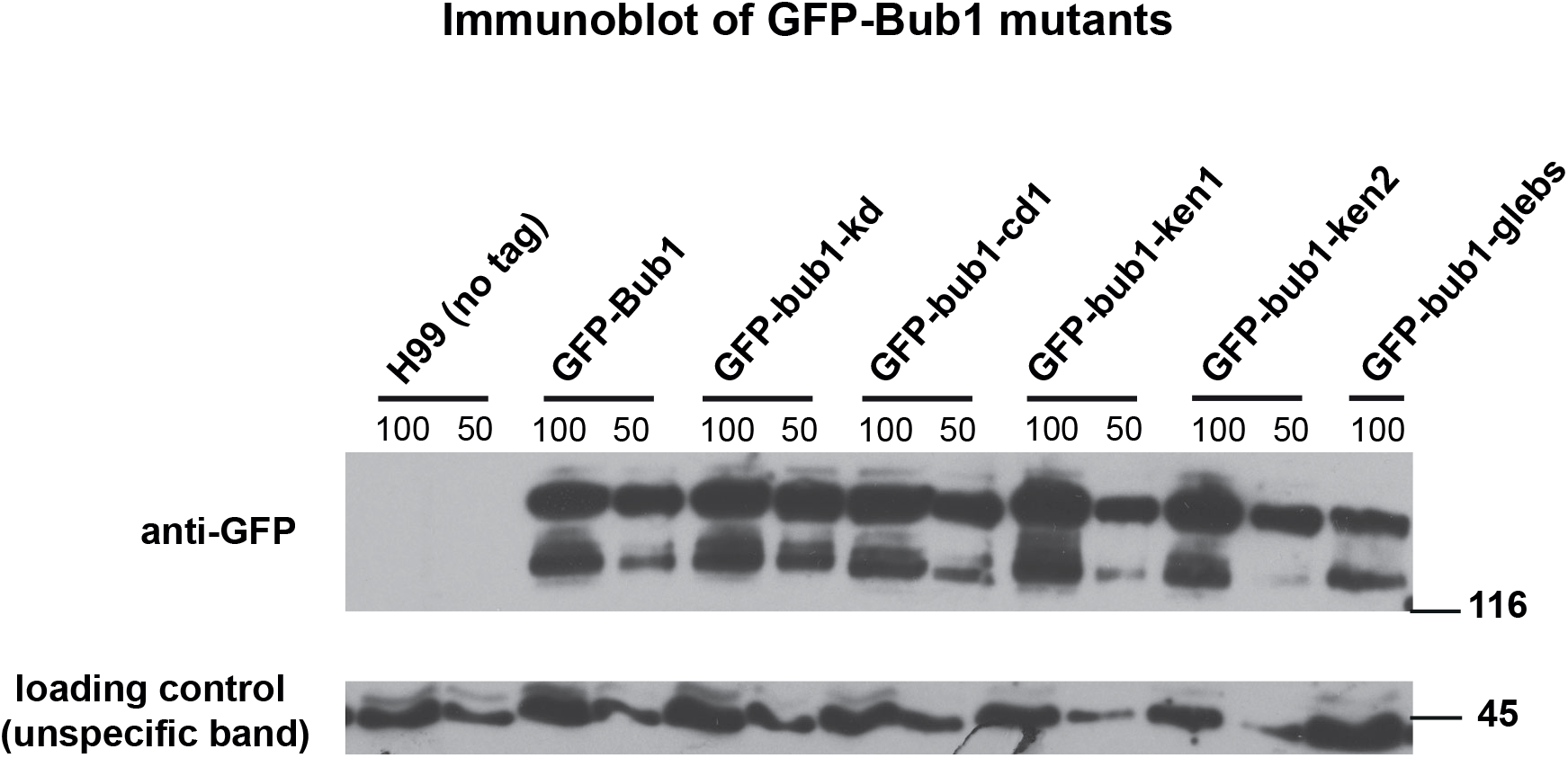
a) Anti-GFP immunoblot of whole cell extracts from H99, *GFP-Bub1* and various *GFP-bub1* mutants. The GFP-bub1-glebs mutant was found at ~50% wildtype levels but other mutant proteins were very similar to wild-type.

**Supp Figure 3.**
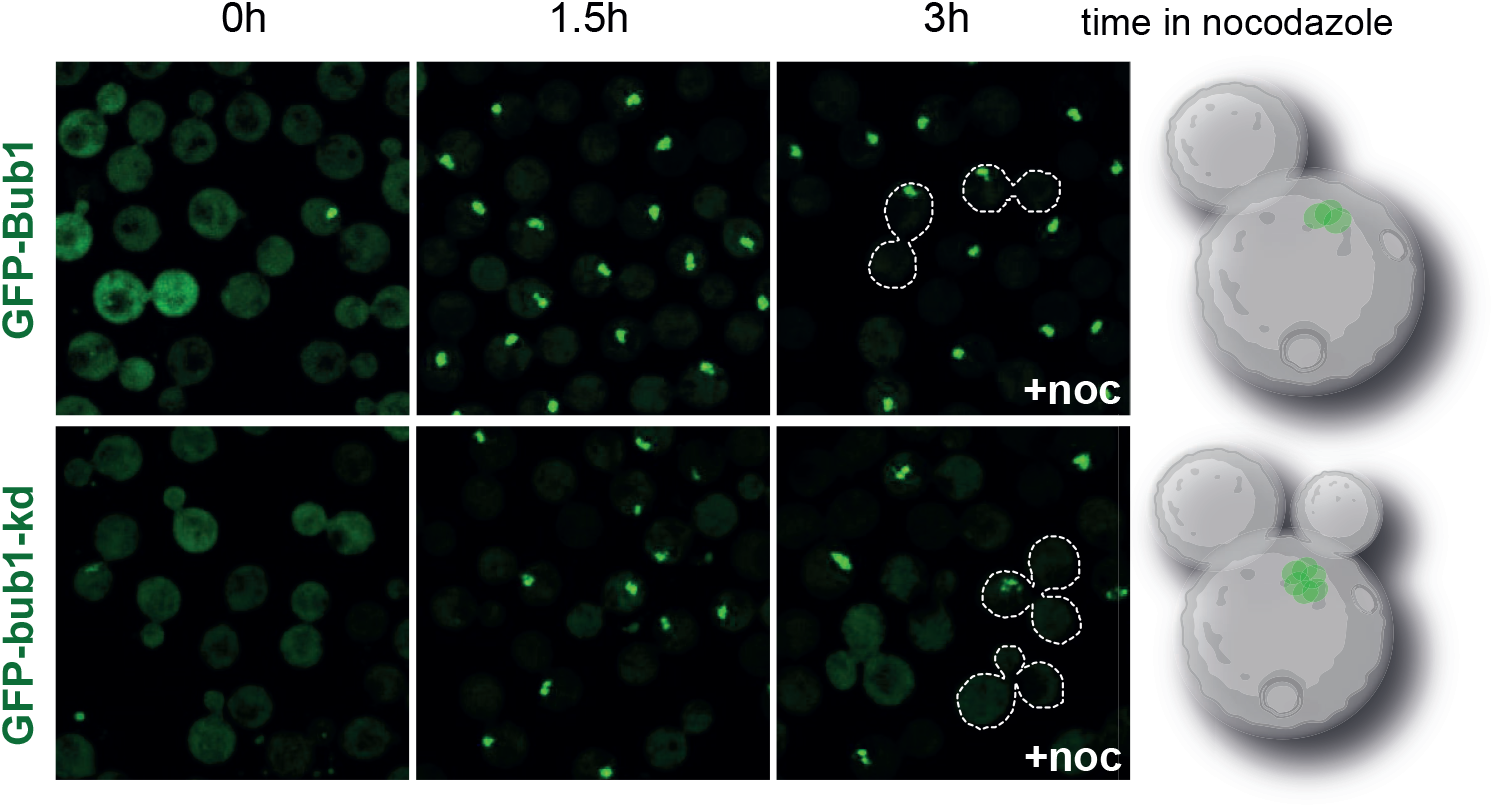
Imaging of the *GFP-Bub1* and *GFP-bub1-kd* strains after 0, 1.5 and 3 hours of nocodazole treatment. The *bub1-kd* cells failed to maintain arrest and re-budded (cells outlined with dashed white lines).

**Supp Figure 4.**
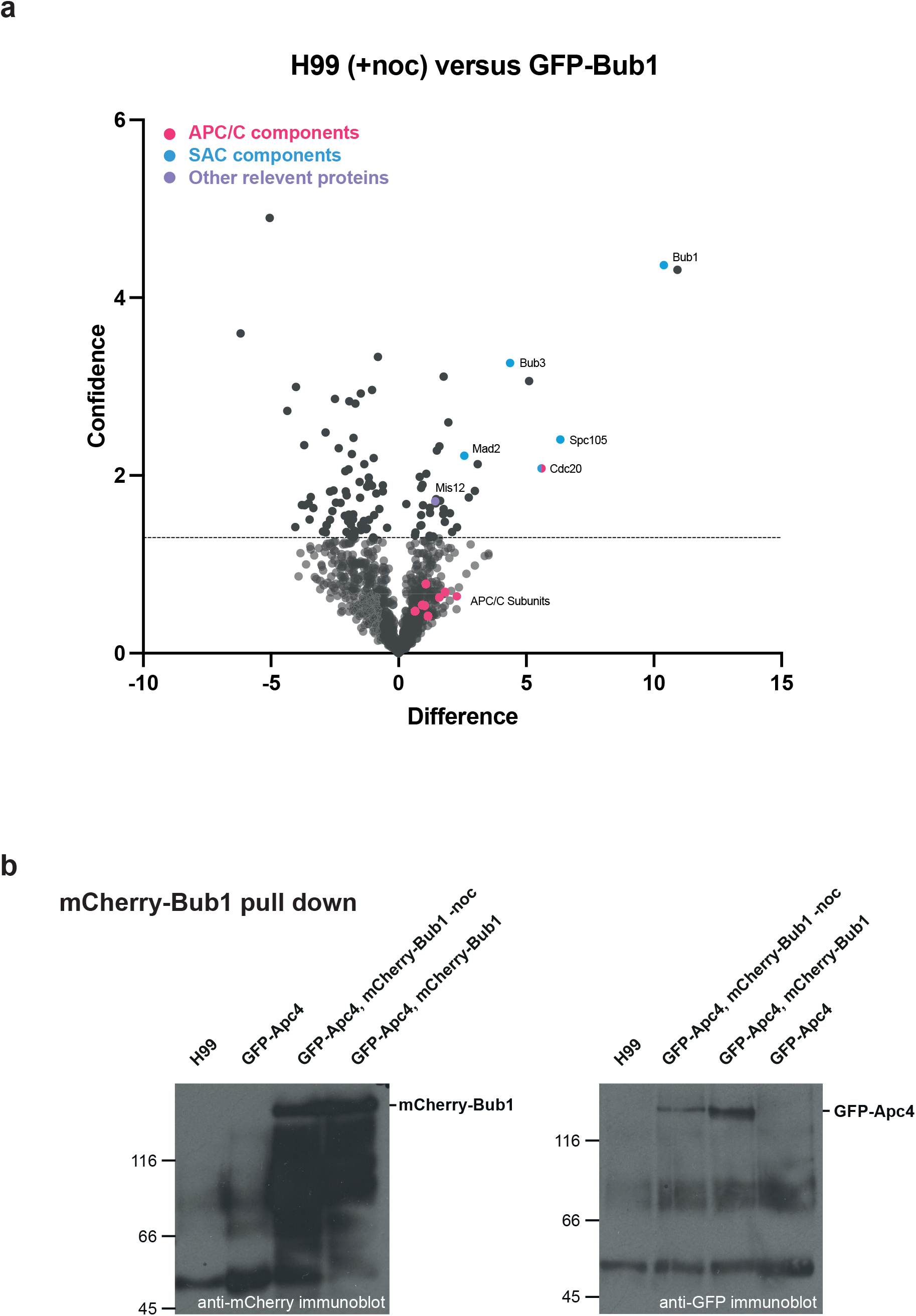
a) Volcano plot comparison of GFP-trap pull downs from extracts of mitotic H99 cells (untagged control) versus cycling GFP-Bub1 cells. b) Anti-mCherry immunoprecipitation pulled down GFP-Apc4 in the strain containing mCherry-Bub1.

## References

1 London, N. & Biggins, S. Signalling dynamics in the spindle checkpoint response. Nat Rev Mol Cell Biol 15, 736–748, doi:nrm3888 [pii] 10.1038/nrm3888 (2014).

2 Kops, G., Snel, B. & Tromer, E. C. Evolutionary Dynamics of the Spindle Assembly Checkpoint in Eukaryotes. Curr Biol 30, R589–R602, doi:10.1016/j.cub.2020.02.021 (2020).

3 Ruchaud, S., Carmena, M. & Earnshaw, W. C. Chromosomal passengers: conducting cell division. Nat Rev Mol Cell Biol 8, 798–812, doi:nrm2257[pii]10.1038/nrm2257 (2007).

4 Roberts, B. T., Farr, K. A. & Hoyt, M. A. The Saccharomyces cerevisiae checkpoint gene BUB1 encodes a novel protein kinase. Mol Cell Biol 14, 8282–8291 (1994).

5 Vleugel, M., Hoogendoorn, E., Snel, B. & Kops, G. J. Evolution and function of the mitotic checkpoint. Dev Cell 23, 239–250, doi:10.1016/j.devcel.2012.06.013 (2012).

6 London, N. & Biggins, S. Mad1 kinetochore recruitment by Mps1-mediated phosphorylation of Bub1 signals the spindle checkpoint. Genes Dev 28, 140–152, doi:gad.233700.113 [pii] 10.1101/gad.233700.113 (2014).

7 Lara-Gonzalez, P., Kim, T., Oegema, K., Corbett, K. & Desai, A. A tripartite mechanism catalyzes Mad2-Cdc20 assembly at unattached kinetochores. Science 371, 64–67, doi:10.1126/science.abc1424 (2021).

8 Piano, V. et al. CDC20 assists its catalytic incorporation in the mitotic checkpoint complex. Science 371, 67–71, doi:10.1126/science.abc1152 (2021).

9 Sudakin, V., Chan, G. K. & Yen, T. J. Checkpoint inhibition of the APC/C in HeLa cells is mediated by a complex of BUBR1, BUB3, CDC20, and MAD2. J Cell Biol 154, 925–936 (2001).

10 Hardwick, K. G., Johnston, R. C., Smith, D. L. & Murray, A. W. MAD3 encodes a novel component of the spindle checkpoint which interacts with Bub3p, Cdc20p, and Mad2p. J Cell Biol 148, 871–882 (2000).

11 Izawa, D. & Pines, J. The mitotic checkpoint complex binds a second CDC20 to inhibit active APC/C. Nature 517, 631–634, doi:nature13911 [pii] 10.1038/nature13911 (2014).

12 Alfieri, C. et al. Molecular basis of APC/C regulation by the spindle assembly checkpoint. Nature 536, 431–436, doi:10.1038/nature19083 (2016).

13 Goldman, D. L. et al. Serologic evidence for Cryptococcus neoformans infection in early childhood. Pediatrics 107, E66, doi:10.1542/peds.107.5.e66 (2001).

14 Rajasingham, R. et al. The global burden of HIV-associated cryptococcal infection in adults in 2020: a modelling analysis. Lancet Infect Dis, doi:10.1016/S1473-3099(22)00499-6 (2022).

15 Zaragoza, O. & Nielsen, K. Titan cells in Cryptococcus neoformans: cells with a giant impact. Curr Opin Microbiol 16, 409–413, doi:10.1016/j.mib.2013.03.006 (2013).

16 Okagaki, L. H. & Nielsen, K. Titan Cells Confer Protection from Phagocytosis in Cryptococcus neoformans Infections. Eukaryot Cell 11, 820–826, doi:10.1128/Ec.00121-12 (2012).

17 Gerstein, A. C. et al. Polyploid Titan Cells Produce Haploid and Aneuploid Progeny To Promote Stress Adaptation. Mbio 6, doi:ARTN e01340-15 10.1128/mBio.01340-15 (2015).

18 Wertheimer, N. B., Stone, N. & Berman, J. Ploidy dynamics and evolvability in fungi. Philos Trans R Soc Lond B Biol Sci 371, doi:10.1098/rstb.2015.0461 (2016).

19 Yang, F. et al. Adaptation to Fluconazole via Aneuploidy Enables Cross-Adaptation to Amphotericin B and Flucytosine in Cryptococcus neoformans. Microbiol Spectr 9, e0072321, doi:10.1128/Spectrum.00723-21 (2021).

20 Janbon, G. et al. Characterizing the role of RNA silencing components in Cryptococcus neoformans. Fungal Genet Biol 47, 1070–1080, doi:10.1016/j.fgb.2010.10.005 (2010).

21 Janbon, G. et al. Analysis of the genome and transcriptome of Cryptococcus neoformans var. grubii reveals complex RNA expression and microevolution leading to virulence attenuation. PLoS Genet 10, e1004261, doi:10.1371/journal.pgen.1004261 (2014).

22 Catania, S. et al. Evolutionary Persistence of DNA Methylation for Millions of Years after Ancient Loss of a De Novo Methyltransferase. Cell 180, 263–277 e220, doi:10.1016/j.cell.2019.12.012 (2020).

23 Kozubowski, L. et al. Ordered kinetochore assembly in the human-pathogenic basidiomycetous yeast Cryptococcus neoformans. Mbio 4, e00614–00613, doi:10.1128/mBio.00614-13 (2013).

24 Sridhar, S., Hori, T., Nakagawa, R., Fukagawa, T. & Sanyal, K. Bridgin connects the outer kinetochore to centromeric chromatin. Nat Commun 12, 146, doi:10.1038/s41467-020-20161-9 (2021).

25 Varshney, N. et al. Spatio-temporal regulation of nuclear division by Aurora B kinase Ipl1 in Cryptococcus neoformans. PLoS Genet 15, e1007959, doi:10.1371/journal.pgen.1007959 (2019).

26 Altamirano, S. et al. The Cyclin Cln1 Controls Polyploid Titan Cell Formation following a Stress-Induced G2 Arrest in Cryptococcus. Mbio 12, e0250921, doi:10.1128/mBio.02509-21 (2021).

27 Lee, K. T. et al. Systematic functional analysis of kinases in the fungal pathogen Cryptococcus neoformans. Nat Commun 7, 12766, doi:10.1038/ncomms12766 (2016).

28 Taylor, S. T., Ha, E. & McKeon, F. The human homolog of Bub3 is required for kinetochore localization of Bub1 and a human Mad3-like protein kinase. Journal of Cell Biology 142, 1–11 (1998).

29 Wickes, B. L. & Edman, J. C. The Cryptococcus neoformans GAL7 gene and its use as an inducible promoter. Mol Microbiol 16, 1099–1109, doi:10.1111/j.1365-2958.1995.tb02335.x (1995).

30 Erpf, P. E., Stephenson, C. J. & Fraser, J. A. amdS as a dominant recyclable marker in Cryptococcus neoformans. Fungal Genet Biol 131, 103241, doi:10.1016/j.fgb.2019.103241 (2019).

31 Murphy, R., Watkins, J. L. & Wente, S. R. GLE2, a *Saccharomyces cerevisiae* homologue of the *Schizosaccharomyce pombe* export factor RAE1, is required for nuclear pore complex structure and function. Molecular Biology of the Cell 7, 1921–1937 (1996).

32 Primorac, I. et al. Bub3 reads phosphorylated MELT repeats to promote spindle assembly checkpoint signaling. Elife 2, e01030, doi:10.7554/eLife.01030 [pii] (2013).

33 London, N., Ceto, S., Ranish, J. A. & Biggins, S. Phosphoregulation of Spc105 by Mps1 and PP1 regulates Bub1 localization to kinetochores. Curr Biol 22, 900–906, doi:10.1016/j.cub.2012.03.052 S0960-9822(12)00339-9 [pii] (2012).

34 Shepperd, L. A. et al. Phosphodependent Recruitment of Bub1 and Bub3 to Spc7/KNL1 by Mph1 Kinase Maintains the Spindle Checkpoint. Curr Biol 22, 891–899, doi:S0960-9822(12)00338-7 [pii] 10.1016/j.cub.2012.03.051 (2012).

35 Yamagishi, Y., Yang, C. H., Tanno, Y. & Watanabe, Y. MPS1/Mph1 phosphorylates the kinetochore protein KNL1/Spc7 to recruit SAC components. Nat Cell Biol 14, 746–752, doi:10.1038/ncb2515 [pii] (2012).

36 Vleugel, M. et al. Sequential multisite phospho-regulation of KNL1-BUB3 interfaces at mitotic kinetochores. Mol Cell 57, 824–835, doi:S1097-2765(14)01014-4 [pii] 10.1016/j.molcel.2014.12.036 (2015).

37 Jenni, S. & Harrison, S. C. Structure of the DASH/Dam1 complex shows its role at the yeast kinetochore-microtubule interface. Science 360, 552–558, doi:10.1126/science.aar6436 (2018).

38 Elowe, S. & Bolanos-Garcia, V. M. The spindle checkpoint proteins BUB1 and BUBR1: (SLiM)ming down to the basics. Trends Biochem Sci 47, 352–366, doi:10.1016/j.tibs.2022.01.004 (2022).

39 Arras, S. D., Chitty, J. L., Blake, K. L., Schulz, B. L. & Fraser, J. A. A genomic safe haven for mutant complementation in Cryptococcus neoformans. PLoS One 10, e0122916, doi:10.1371/journal.pone.0122916 (2015).

40 Heinrich, S. et al. Mad1 contribution to spindle assembly checkpoint signalling goes beyond presenting Mad2 at kinetochores. EMBO Rep 15, 291–298, doi:10.1002/embr.201338114 (2014).

41 King, E. M., van der Sar, S. J. & Hardwick, K. G. Mad3 KEN boxes mediate both Cdc20 and Mad3 turnover, and are critical for the spindle checkpoint. PLoS ONE 2, e342 (2007).

42 Lara-Gonzalez, P., Scott, M. I., Diez, M., Sen, O. & Taylor, S. S. BubR1 blocks substrate recruitment to the APC/C in a KEN-box-dependent manner. J Cell Sci 124, 4332–4345, doi:jcs.094763 [pii] 10.1242/jcs.094763 (2011).

43 Di Fiore, B. et al. The ABBA motif binds APC/C activators and is shared by APC/C substrates and regulators. Dev Cell 32, 358–372, doi:10.1016/j.devcel.2015.01.003 (2015).

44 Brady, D. M. & Hardwick, K. G. Complex formation between Mad1p, Bub1p and Bub3p is crucial for spindle checkpoint function. Current Biology 10, 675–678 (2000).

45 Fischer, E. S. et al. Molecular mechanism of Mad1 kinetochore targeting by phosphorylated Bub1. EMBO Rep 22, e52242, doi:10.15252/embr.202052242 (2021).

46 Aktar, K., Leontiou, I. & Hardwick, K. G. in preparation.

47 Meraldi, P. Bub1-the zombie protein that CRISPR cannot kill. EMBO J 38, doi:10.15252/embj.2019101912 (2019).

48 Raaijmakers, J. A. & Medema, R. H. Killing a zombie: a full deletion of the BUB1 gene in HAP1 cells. EMBO J 38, e102423, doi:10.15252/embj.2019102423 (2019).

49 Crane, M. M., Clark, I. B., Bakker, E., Smith, S. & Swain, P. S. A microfluidic system for studying ageing and dynamic single-cell responses in budding yeast. PLoS One 9, e100042, doi:10.1371/journal.pone.0100042 (2014).

50 Sewart, K. & Hauf, S. Different functionality of Cdc20 binding sites within the mitotic checkpoint complex. (2017).

51 May, K. M., Paldi, F. & Hardwick, K. G. Fission Yeast Apc15 Stabilizes MCC-Cdc20-APC/C Complexes, Ensuring Efficient Cdc20 Ubiquitination and Checkpoint Arrest. Curr Biol 27, 1221–1228, doi:10.1016/j.cub.2017.03.013 (2017).

52 Sczaniecka, M. et al. The spindle checkpoint functions of Mad3 and Mad2 depend on a Mad3 KEN box-mediated interaction with Cdc20-anaphase-promoting complex (APC/C). J Biol Chem 283, 23039–23047 (2008).

53 Kawashima, S. A., Yamagishi, Y., Honda, T., Ishiguro, K. & Watanabe, Y. Phosphorylation of H2A by Bub1 prevents chromosomal instability through localizing shugoshin. Science 327, 172–177, doi:science.1180189 [pii] 10.1126/science.1180189 (2010).

54 Tang, Z., Shu, H., Oncel, D., Chen, S. & Yu, H. Phosphorylation of Cdc20 by Bub1 provides a catalytic mechanism for APC/C inhibition by the spindle checkpoint. Mol Cell 16, 387–397 (2004).

55 Jia, L., Li, B. & Yu, H. The Bub1-Plk1 kinase complex promotes spindle checkpoint signalling through Cdc20 phosphorylation. Nat Commun 7, 10818, doi:10.1038/ncomms10818 (2016).

56 Shah, J. V. et al. Dynamics of centromere and kinetochore proteins; implications for checkpoint signaling and silencing. Curr Biol 14, 942–952 (2004).

57 Howell, B. J. et al. Spindle checkpoint protein dynamics at kinetochores in living cells. Curr Biol 14, 953–964 (2004).

58 Nguyen Ba, A. N. et al. Parallel reorganization of protein function in the spindle checkpoint pathway through evolutionary paths in the fitness landscape that appear neutral in laboratory experiments. PLoS Genet 13, e1006735, doi:10.1371/journal.pgen.1006735 (2017).

59 Kawashima, S. A. et al. Shugoshin enables tension-generating attachment of kinetochores by loading Aurora to centromeres. Genes Dev 21, 420–435 (2007).

60 Fernius, J. & Hardwick, K. G. Bub1 kinase targets Sgo1 to ensure efficient chromosome biorientation in budding yeast mitosis. PLoS Genet 3, e213 (2007).

61 Hanks, S. et al. Comparative genomic hybridization and BUB1B mutation analyses in childhood cancers associated with mosaic variegated aneuploidy syndrome. Cancer Lett 239, 234–238 (2006).

62 Orner, E. P. et al. High-Throughput Yeast Aging Analysis for Cryptococcus (HYAAC) microfluidic device streamlines aging studies in Cryptococcus neoformans. Commun Biol 2, 256, doi:10.1038/s42003-019-0504-5 (2019).

63 Granados, A. A. et al. Distributed and dynamic intracellular organization of extracellular information. Proc Natl Acad Sci U S A 115, 6088–6093, doi:10.1073/pnas.1716659115 (2018).

64 Granados, A. A. et al. Distributing tasks via multiple input pathways increases cellular survival in stress. Elife 6, doi:10.7554/eLife.21415 (2017).

65 Cai, L., Dalal, C. K. & Elowitz, M. B. Frequency-modulated nuclear localization bursts coordinate gene regulation. Nature 455, 485–490, doi:10.1038/nature07292 (2008).

66 Hao, N. & O’Shea, E. K. Signal-dependent dynamics of transcription factor translocation controls gene expression. Nat Struct Mol Biol 19, 31–39, doi:10.1038/nsmb.2192 (2011).

67 Lin, Y., Sohn, C. H., Dalal, C. K., Cai, L. & Elowitz, M. B. Combinatorial gene regulation by modulation of relative pulse timing. Nature 527, 54–58, doi:10.1038/nature15710 (2015).

68 Abad, M. A. et al. Borealin-nucleosome interaction secures chromosome association of the chromosomal passenger complex. J Cell Biol 218, 3912–3925, doi:10.1083/jcb.201905040 (2019).

69 Shevchenko, A., Wilm, M., Vorm, O. & Mann, M. Mass spectrometric sequencing of proteins silver-stained polyacrylamide gels. Anal Chem 68, 850–858, doi:10.1021/ac950914h (1996).

70 Rappsilber, J., Mann, M. & Ishihama, Y. Protocol for micro-purification, enrichment, pre-fractionation and storage of peptides for proteomics using StageTips. Nat Protoc 2, 1896–1906, doi:nprot.2007.261 [pii] 10.1038/nprot.2007.261 (2007).

71 Cox, J. & Mann, M. MaxQuant enables high peptide identification rates, individualized p.p.b.-range mass accuracies and proteome-wide protein quantification. Nat Biotechnol 26, 1367–1372, doi:10.1038/nbt.1511 (2008).

72 Cox, J. et al. Andromeda: a peptide search engine integrated into the MaxQuant environment. J Proteome Res 10, 1794–1805, doi:10.1021/pr101065j (2011).

73 Cox, J. et al. Accurate proteome-wide label-free quantification by delayed normalization and maximal peptide ratio extraction, termed MaxLFQ. Mol Cell Proteomics 13, 2513–2526, doi:10.1074/mcp.M113.031591 (2014).

74 Tyanova, S. et al. The Perseus computational platform for comprehensive analysis of (prote)omics data. Nat Methods 13, 731–740, doi:10.1038/nmeth.3901 (2016).

